# A single nucleotide change underlies the genetic assimilation of a plastic trait

**DOI:** 10.1101/2020.06.29.176990

**Authors:** Paul Vigne, Clotilde Gimond, Céline Ferrari, Anne Vielle, Johan Hallin, Ania Pino-Querido, Sonia El Mouridi, Christian Frøkjær-Jensen, Thomas Boulin, Henrique Teotónio, Christian Braendle

**Author notes:** equal contribution (Paul Vigne, Clotilde Gimond).

## Abstract

Genetic assimilation – the evolutionary process by which an ancestral environmentally sensitive phenotype is made constitutive – is a fundamental concept in biology. Its evolutionary relevance is debated, and our understanding of its prevalence, and underlying genetics and molecular mechanisms, is poor. Matricidal hatching is an extreme form of maternal provisioning induced by adverse conditions, which varies among *Caenorhabditis elegans* populations. We identified wild isolates, sampled from natural populations across multiple years and locations, that express a derived state of near-constitutive matricidal hatching. A single amino acid change in *kcnl-1*, encoding a small-conductance calcium-activated potassium channel subunit, explains most of this variation. A gain-of-function mutation altering the S6 transmembrane domain causes inappropriate activation of the K^+^ channel, leading to reduced vulval muscle excitability, and thus reduced expulsion of embryos, irrespective of environment. Using reciprocal allelic replacements, we show that this amino acid change is sufficient to induce constitutive matricidal hatching whilst re-establishing the ancestral protein abolishes matricidal hatching and restores egg-laying, thereby doubling lifetime reproductive fitness under benign conditions. While highly deleterious in the laboratory, experimental evolution showed that KNCL-1(V530L) is maintained under fluctuating resource availability. Selection on a single point mutation can therefore underlie the genetic assimilation of an ancestrally plastic trait with drastic life-history consequences.

---

Adaptive phenotypic plasticity is a central organismal strategy to cope with spatio-temporal variation in the environment, allowing a single genotype to express different phenotypes (Bradshaw, 1965; Sommer, 2020; West-Eberhard, 2003). A long-standing debate in biology surrounds the question of how an ancestrally plastic, environmentally-sensitive phenotype can evolve into a fixed, genetically encoded phenotype (Baldwin, 1896; Morgan, 1896; Waddington, 1942). This evolutionary process is commonly referred to as *genetic assimilation*, a term coined by Conrad Hal Waddington (Waddington, 1942). Genetic assimilation received considerable attention in recent years in the context of research focusing on the role of phenotypic plasticity in the evolutionary process (Ghalambor et al., 2007; Lande, 2009; Levis and Pfennig, 2019; Moczek et al., 2011; Pigliucci et al., 2006; Sommer, 2020; Suzuki and Nijhout, 2006; West-Eberhard, 2003). Nevertheless, the molecular mechanisms underlying evolutionary transitions from plastic to fixed trait expression remain poorly understood. Currently we cannot resolve to what extent evolutionary loss of plasticity, leading to fixation of one of several alternative phenotypes, occurs through gradual canalizing selection on environmentally-released cryptic genetic variation (Loison, 2019; Suzuki and Nijhout, 2006; Waddington, 1942, 1959) or through different mechanisms, such as *de novo* mutations. In addition, virtually all insights into the mechanistic basis of genetic assimilation in animals rely on artificial selection experiments (Fanti et al., 2017; Gibson and Hogness, 1996; Suzuki and Nijhout, 2006). Therefore, whether and how genetic assimilation occurs in natural populations remains unclear.

Here we identified the molecular basis underlying the evolutionary transition from an ancestrally plastic to a genetically fixed trait focusing on matricidal hatching in natural populations of the hermaphroditic nematode *Caenorhabditis elegans*. Egg-laying behaviour of *C. elegans* is plastic and sensitive to diverse environmental factors (Horvitz et al., 1982; Schafer, 2006). Self- and cross-fertilization in *C. elegans* is internal, with embryos initially developing for 2-3 hours in the hermaphrodite uterus, and completing embryogenesis 10-12 hours later, under benign environmental conditions (Sulston et al., 1983). Upon starvation, however, hermaphrodites will retain embryos in the uterus where they will hatch into larvae that will continue to grow (“bag of worms” or “facultative viviparity”) and eventually lead to the rapid and premature death of their mother (Chen and Caswell-Chen, 2003; Trent, 1982). In nutrient-scarce environments, facultative matricidal hatching may be an adaptive strategy since mothers provide essential resources for offspring survival by allowing them, for example, to reach the alternative diapausing dauer larval stage (Chen and Caswell-Chen, 2003, 2004). The dauer stage is resistant to nutrient-deprivation and will disperse to colonize novel habitats (Frézal and Félix, 2015). In addition, when facing starvation, early larvae can diapause (Baugh, 2013), and so matricidal hatching might also be adaptive because it protects the developing non-diapausing offspring whom will head-start population growth when conditions become favourable. Hence, matricidal hatching is thought to be an extreme form of offspring provisioning or protection, even at the cost of maternal survival, that is favored by natural selection when populations face fluctuating environments across generations (Dey et al., 2016; Proulx and Teotónio, 2017).

Plasticity in *C. elegans* matricidal hatching is conserved across genetically divergent wild isolates (Fig. 1A) (Chen and Caswell-Chen, 2004; Stastna et al., 2015), with the notable exception of the isolate JU751, which displays a constitutively high rate of matricidal hatching irrespective of environmental variation (Figs. 1A and 1B). At the phenotypic level, fixed matricidal hatching JU751 thus appears to be derived from an ancestrally plastic state. Although JU751 hermaphrodites laid eggs during early adulthood, they exhibited internally hatched L1 larvae later in life (Fig. 1B). Comparing JU751 to the wild isolate JU1200 with ancestral *C. elegans* egg-laying behaviour in an *ad libitum* food environment, we found JU751 to have significantly increased egg retention and correspondingly high matricidal hatching (Figs. 1C to 1F and figs. S1 and S2). Internal hatching led to early maternal death (Figs. 1G, S3A and S3B), because larvae moved vigorously inside the mother causing extensive injury to somatic and gonadal tissues (Fig. S2). Internal larval hatching of JU751 individuals in benign conditions thus seems to carry a considerable fitness cost because maternal death occurred before the end of the reproductive span, prior to exhaustion of hermaphrodite self-sperm (Figs. S3C and S3D).

**Fig. 1.**
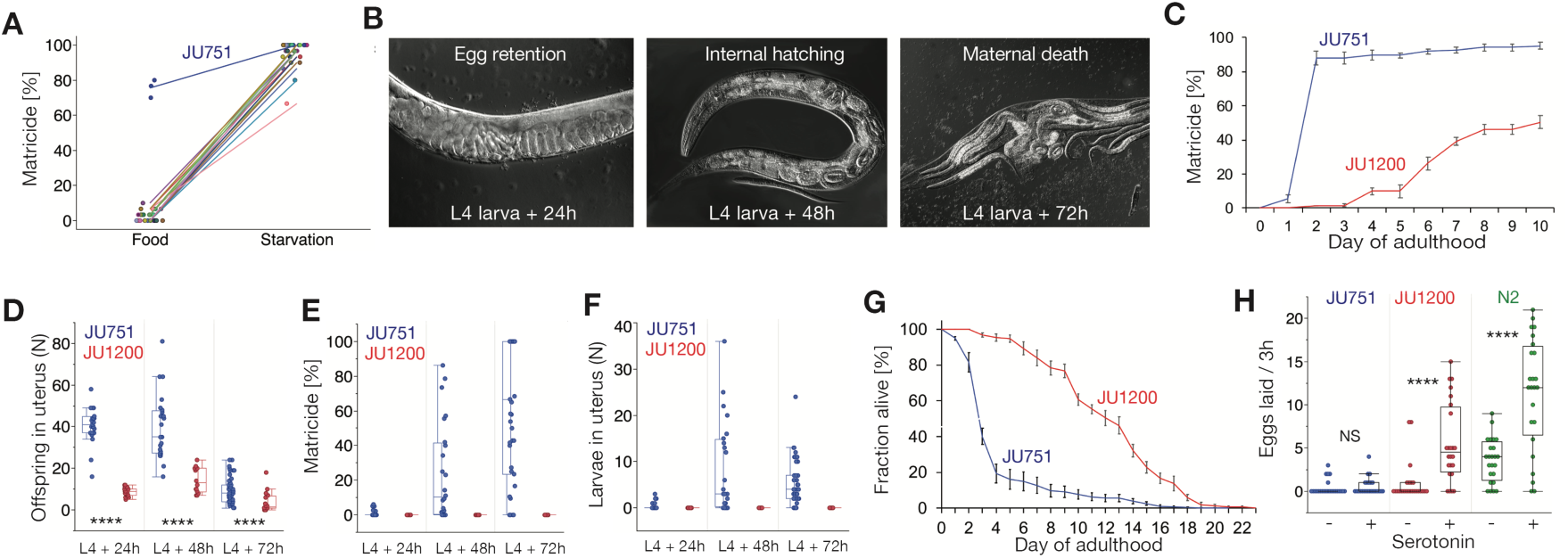
Natural variation in plasticity of *C. elegans* matricidal hatching. (**A**) Natural variation in the plasticity of matricidal hatching across genetically divergent *C. elegans* wild isolates (N=28) quantified in food (solid) and starvation (liquid) culture. Values of matricidal hatching reflect the percentage of mothers containing one or more internally hatched larvae (N=30 individuals/strain/environment; except for JU751: N=3×30 individuals). (**B**) Temporal progression of matricidal hatching of JU751 hermaphrodites in *ad libitum* food conditions (Nomarski micrographs). (**C**) Temporal dynamics of matricidal hatching in JU751 versus JU1200: cumulative percentage of individuals containing one or more internally hatched larvae (N=5 replicates/strain, N=29-32 individuals/replicate). For further details, see figs. S1C to S1E. (**D**-**F**) Egg retention and internal hatching in JU751 and JU1200 in *ad libitum* food conditions at 20°C (N=14-47 individuals/strain/time point) (see also fig. S1B). (**D**) Number of offspring retained *in utero* (embryos and larvae) during the reproductive span. (ANOVA performed separately for each time point: \*\*\*\**P* < 0.0001). (**E**) Proportion of mothers displaying matricidal hatching, i.e. mothers containing one or more internally hatched larva. (**F**) Total number of internally developing larvae. (**G**) Survival curves of JU751 and JU1200 hermaphrodites in *ad libitum* food conditions at 20°C (N=9-15 replicates, N=9-44 individuals/replicate). Matricidal hatching prior to day five of adulthood accounts for >60% of mortality in JU751. For further details, see figs. S3A and S3B. (**H**) Exogenous application of serotonin (5-HT) triggered egg-laying in JU1200 and the reference strain N2 but not in JU751 (liquid starvation culture, N=24 individuals/strain/treatment) (ANOVA performed separately for each strain: NS, not significant, \*\*\*\**P* < 0.0001).

Multiple observations suggest that the egg-laying apparatus of JU751 is not simply defective. First, JU751 hermaphrodites laid eggs containing advanced embryos during early adulthood (Fig. S1B). Second, egg-laying activity in young adults was the same in JU751 and JU1200, with similar temporal patterns of active and inactive egg-laying phases (Figs. S4C and S4D), although JU751 laid fewer eggs per active event (Figs. S4E and S4F). Third, we did not detect any obvious defects in vulval development, feeding (pumping), food sensory behaviour (olfaction), or morphology that could be indicative of starvation when fed on the standard *E. coli* food source. Finally, egg retention and matricidal hatching of JU751 were also similar and consistently high across different food sources or temperatures (Figs. S1C to S1E). Therefore, the state of constitutive matricidal hatching in JU751 is consistent with a scenario of reduction in environmental sensitivity of the egg-laying circuit. This interpretation is supported by JU751’s lack of response to the neurotransmitter serotonin – known to mimic food-stimulation of *C. elegans* egg-laying (Horvitz et al., 1982; Schafer, 2006; Trent, 1982) (Fig. 1H).

To characterize the genetic basis of constitutive matricidal hatching in JU751, we derived F2 Recombinant Inbred Lines (RILs) from a reciprocal parental cross between isolates JU751 and JU1200 (Figs. 2A and S5A). 144 SNP-genotyped RILs (Fig. S6) were scored for the presence or absence of internally hatched larvae within the first three days of adulthood at 15°C (Figs. 2B and S5B). Quantitative Trait Locus (QTL) mapping uncovered a single significant QTL on chromosome V, spanning approximately 1Mb (Fig. 2C). The QTL was validated by generating Near-Isogenic Lines (NILs) carrying the QTL region with JU751 genotype backcrossed into the JU1200 background: all NILs carrying the introgressed QTL region showed strong egg retention and matricidal hatching (Fig. S7A). Additional genotyping of the QTL region restricted the target interval to 60kb (Fig. S7B) covering three protein-coding genes, which contained a single polymorphism, located in the coding region of the gene *kcnl-1* (B0399.1) (Figs. 2D and S7B).

**Fig. 2.**
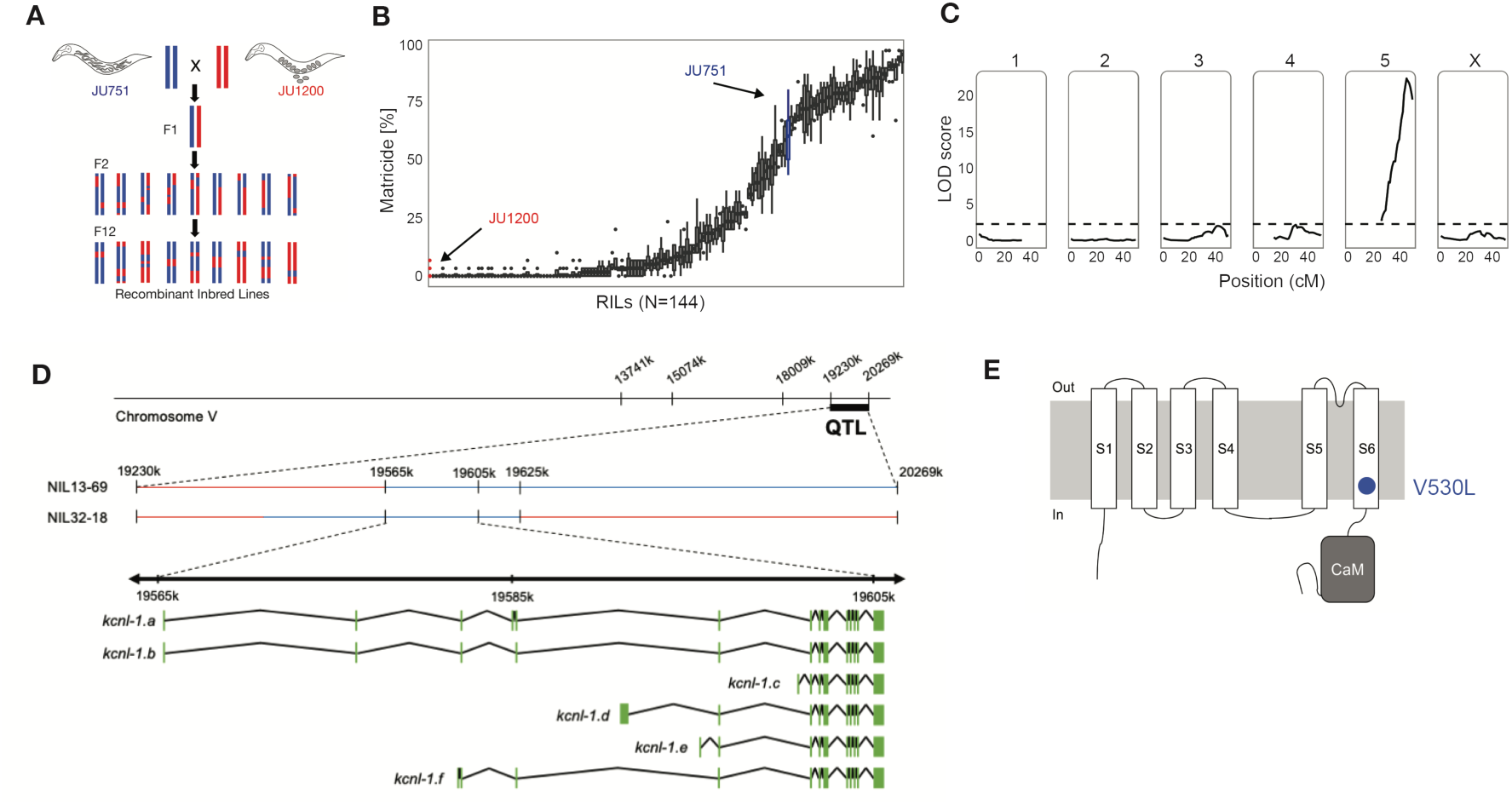
Quantitative Trait Locus (QTL) mapping identifies a single large-effect locus explaining variation in matricidal hatching. (**A**) Crossing scheme to generate F2 RILs derived from a parental cross between isolates JU751 and JU1200 (for details, see Fig. S5A). (**B**) Phenotypic distribution (cumulative percentage of individuals undergoing matricidal hatching at mid-L4+72h at 15°C) of all replicates (N=4-5) for each RIL (N=144). Each boxplot displays all replicate values for a given RIL (*x*-axis). Parental strains are coloured in red (JU1200) and blue (JU751). (**C**) QTL map of the LOD scores for the percentage of matricidal hatching at mid-L4+72 hours. The dashed horizontal line shows the 5% significance threshold calculated with 1000 permutations. Phenotype data from 144 RILs was used for the mapping in R/qtl. (**D**) From QTL to QTN: fine-mapping through near-isogenic lines and additional genotyping of the target region identifies a single polymorphism in the gene *kncl-1*, a small conductance (SK) calcium-activated potassium channel subunit with six predicted isoforms, ranging from 2606 (isoform a) to 1937 (isoform c) amino acids. (**E**) Predicted structure of the KCNL-1 ion channel subunit (Köhler et al., 1996). The KCNL-1(V530L) variant in JU751 affects the predicted S6 transmembrane (TM) domain.

*kcnl-1* encodes a small conductance (SK) calcium-activated potassium channel subunit with six predicted isoforms (Fig. 2D). KCNL-1 is one of four *C. elegans* orthologues of the human *KCNN* SK potassium channel family, characterized by six transmembrane segments and a C-terminal Calmodulin (CaM) binding domain (Fig. 2E). The JU751 *kcnl-1* variant is a single nucleotide change (**G**TG **→ T**TG) causing a valine to leucine substitution (V530L) in the region encoding the S6 transmembrane segment (TM S6) (Fig. 2E). The KCNL-1 protein, including the TM S6 region, shows substantial divergence across distant taxa (Fig. S8A). However, the KCNL-1 V530L variant in JU751 seems exceptional as the affected TM S6 region is highly conserved, not only across all examined (>30) species of the genus *Caenorhabditis*, but also across diverse nematode taxa encompassing hundreds of millions of years of species divergence (Blaxter et al., 1998; Kiontke and Fitch, 2005) (Fig. S8B).

We characterized an additional *kcnl-1* mutation by cloning *exp-3(n2372)*, initially isolated as a semi-dominant mutation that disrupts the expulsion step of the defecation cycle (Reiner et al., 1995) and increases egg retention. This mutation causes a single amino acid change in KCNL-1 (A443V), situated in the intracellular loop that connects the S4 and S5 transmembrane segments of KCNL-1 (Fig. S9A). Both *exp-3(n2372)* and JU751 showed high egg retention (Fig. S9B) and an increased defecation cycle length (Fig. S9C). Egg-laying defects of *exp-3(n2372)* could be suppressed by RNAi (Fig. S9D) and a suppressor screen uncovered only intragenic loss-of-function mutations (unpublished observation). Furthermore, overexpression of mutant *kcnl-1(A443V)* cDNA was sufficient to cause egg retention (Fig. S9E) strongly arguing for a gain-of-function effect of the A443V amino acid change. In contrast, and as expected (Salkoff et al., 2005), a presumptive loss-of-function (deletion) allele of *kcnl-1* was similar to wild type, causing only a slight reduction in egg retention (Fig. S9B).

To identify the focus of action of *kcnl-1*, we investigated its expression pattern by generating a transcriptional reporter strain using CRISPR-*Cas9* gene editing (Figs. 3A to 3D). Consistent with the phenotype of *kcnl-1* gain-of-function mutants, we observed strong expression in neurons and muscles involved in egg-laying (Fig. 3A) and defecation (Fig. 3B). In addition, *kcnl-1* was expressed in body-wall muscles, a few neurons in head and tail ganglia, in ventral nerve cord motoneurons, and in the PVD mechanosensory neuron (Figs. 3C and 3D). To discriminate between a role of KCNL-1 in vulval muscles versus the cholinergic VC neurons that innervate them, we restricted expression of the KCNL-1 V530L mutant to vulval muscles by using a *kcnl-1* promoter fragment that drives expression in vulval muscles but not VC neurons. Mosaic analysis showed that animals carrying this transgene in vulval muscles were incapable of laying eggs (N=492±5), while mosaic animals lacking the transgene in vulval muscles could lay eggs normally (N=34) (Fig. S9F). The ability of RNAi, which is inefficient in neurons, to suppress the egg-laying phenotype (Fig. S9D) further supported a non-neuronal role for KNCL-1. Matricidal hatching in KCNL-1 gain-of-function variants thus likely results from a hyperpolarization of vulval muscles that leads to disruption of egg laying.

**Fig. 3.**
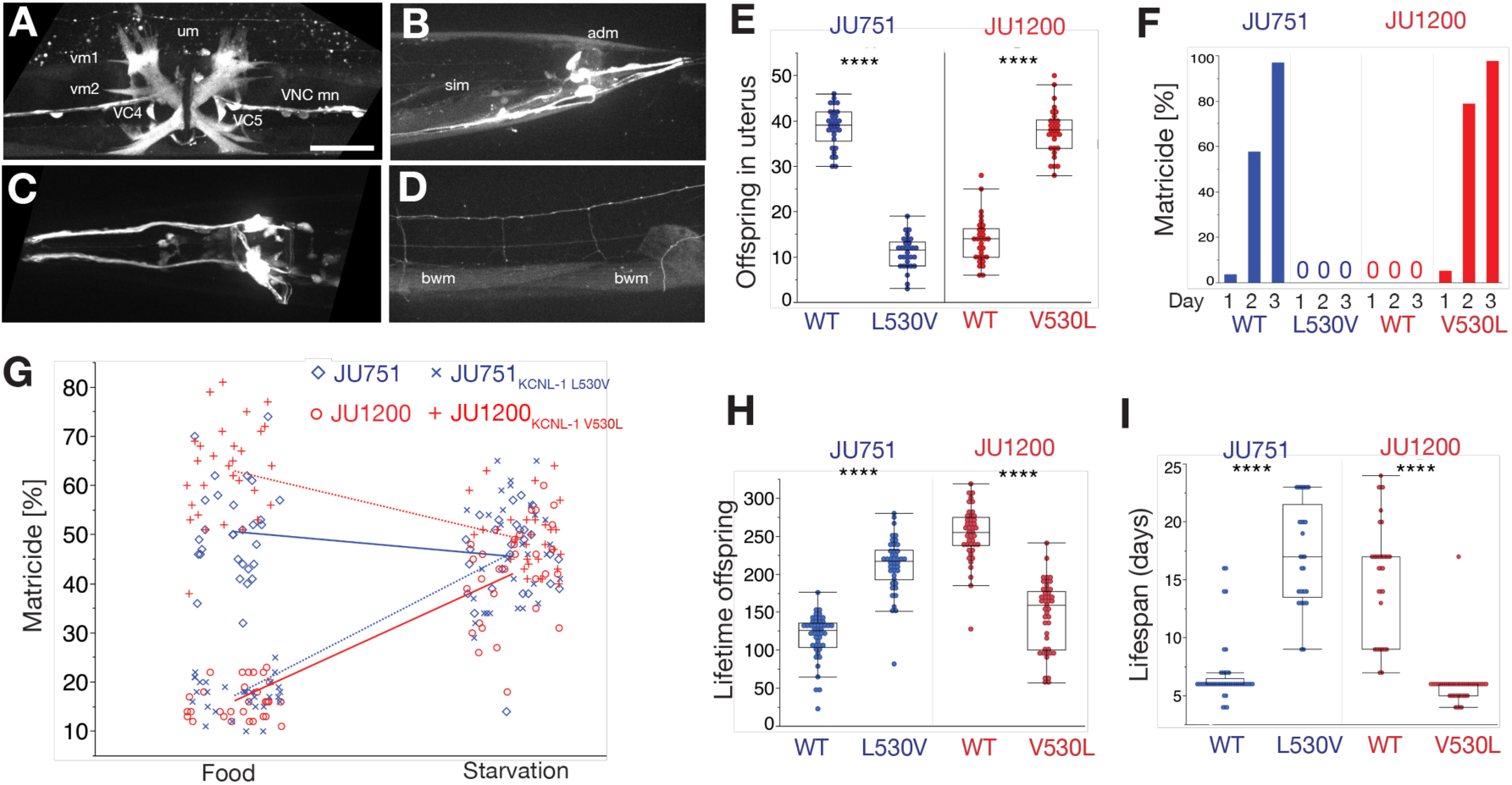
A single amino acid change in KCNL-1 causes constitutive matricidal hatching in JU751. (**A**-**D**) Expression analysis *kcnl-1(bln508[kcnl-1::SL2::wrmScarlet])*. Scale bar, 20μm. (**A**) Strong expression in vulval muscle 1 (vm1) and 2 (vm2), uterine muscle (um), ventral nerve chord motoneurons (VNC mn), and VC4/5 cholinergic neurons. (**B**) Expression in somato-intestinal muscle (sim) and anal depressor muscle (adm). (**C**) Expression in neurons of the head and tail ganglia, in the ventral nerve cord motoneurons. (**D**) Weak expression in body-wall muscles (bwm). (**E**) Offspring number in uterus (embryos and larvae) of adult hermaphrodites (L4+48h) in JU751_WT_, JU751 ARL_KCNL-1 L530V_ and JU1200_WT_, JU1200 ARL_KCNL-1 V530L._ N=30/strain. (**F**) Temporal dynamics of matricidal hatching in JU751_WT_, JU751 ARL_KCNL-1 L530V_ and JU1200_WT_, JU1200 ARL_KCNL-1 V530L_ across the first three days of adulthood (L4+24h, L4+48h, L4+72h). Data show cumulative percentage of mothers displaying matricidal hatching, i.e. mothers containing one or more internally hatched larva. N=119-140/strain. (**G**) Offspring number in uterus (embryos and larvae) of adult hermaphrodites (L4+36h) in JU751_WT_, JU751 ARL_KCNL-1 L530V_ and JU1200_WT_, JU1200 ARL_KCNL-1 V530L_ in food (solid) versus starvation (liquid). N=27-30/strain. (2-Way ANOVA, *genotype*: F_3,229_=176.1, P<0.0001; *environment*: F_1,229_=132.4, P<0.0001; *genotype* x *environment*: F_3,229_=124.5, P<0.0001). (**H**) Lifetime offspring production of selfing hermaphrodites in JU751_WT_, JU751 ARL_KCNL-1 L530V_ and JU1200_WT_, JU1200 ARL_KCNL-1 V530L_. N=43-47/strain. (**I**) Lifespan of selfing hermaphrodites in JU751_WT_, JU751 ARL_KCNL-1 L530V_ and JU1200_WT_, JU1200 ARL_KCNL-1 V530L_. N=29-45/strain. (Figs. **E, H** and **I:** ANOVAs performed separately for each strain background: NS, not significant, \**P* < 0.05, \*\**P* < 0.01, \*\*\**P* < 0.001, \*\*\*\**P* < 0.0001).

We next constructed reciprocal allelic replacement lines (ARLs) in JU751 and JU1200 using CRISPR-*Cas9* gene editing by targeting the underlying single-nucleotide variant (**G**TG **→ T**TG). The resulting amino acid change in the JU1200 allelic replacement line (ARL_KCNL-1 V530L_) led to strong egg retention and highly penetrant matricidal hatching whilst the reciprocal nucleotide replacement in JU751 (ARL_KCNL-1_ L530V) restored fully functional egg-laying and abolished matricidal hatching (Figs. 3E, 3F and S9G). Moreover, introducing the canonical KCNL-1 protein into JU751 also re-established the ancestral state of strong environment-dependent plasticity of *C. elegans* matricidal hatching (Fig. 3G). Hence, KCNL-1 V530L represents the central molecular change underlying the evolutionary transition from ancestrally plastic to constitutive matricidal hatching in the *C. elegans* wild isolate JU751. Measuring life history traits in allelic replacement lines confirms the highly deleterious nature of the KCNL-1 V530L variant: re-establishing the canonical KCNL-1 protein in JU751 nearly doubled its lifetime reproductive output (Fig. 3H) and tripled lifespan (Fig. 3I).

The presence of the *kcnl-1* gain-of-function allele in JU751 with strongly deleterious effects raised the question of whether this variant is truly of natural origin. We characterized independently isolated strains from the JU751 natural habitat, a compost heap in an urban garden close to Paris (France), which was repeatedly sampled during 2004 and 2005 (Barrière and Félix, 2007) (Figs. S10A and S10B). Out of 36 isolates, five additional isolates (isolated on the same day in June of 2005) exhibited the KCNL-1 V530L variant, which displayed strong egg retention and matricidal hatching as in JU751 (Fig. S10C), consistent with a natural origin of this variant. Exploring whole-genome sequence variant data of 249 *C. elegans* wild isotypes (Cook et al., 2017), we also detected coding sequence variants in additional wild isolates (Fig. S11A), including KCNL-1 V530L, which was present in two wild isolates collected from garden compost: JU2587 (Haute Loire, France, 2013) and JU2593 (Hauts-de-Seine, France, 2013) (Figs. S11B and S11C). These isolates also displayed strong egg retention and matricidal hatching (Fig. S11D) and are closely related to JU751 (Fig. S11B) based on analysis of whole-genome sequence similarity(Cook et al., 2017), suggestive of a single evolutionary origin of KCNL-1 V530L. The presence of the KCNL-1 V530L variant in distinct isolates from different localities, sampled across a span of eight years, suggests that this variant is being maintained in natural *C. elegans* populations.

How can the KCNL-1 V530L variant be maintained in natural populations given its seemingly strong deleterious effects as assessed in standard laboratory environment with *ad libitum* food? Given that mutational effects may be strongly dependent on environmental factors, one possibility is that these deleterious effects of KCNL-1 V530L are alleviated in less benign environments. We mimicked more natural conditions of the *C. elegans* habitat (Frézal and Félix, 2015) by exposing JU751, JU1200 and their corresponding *kcnl-1* allelic replacement lines to starvation (liquid culture) at different maternal ages. Stress exposure experienced at multiple time points during the reproductive span strongly reduced, or eliminated, the negative fitness effects of KCNL-1 V530L (Figs. 4A and S12A). Introducing this variant into JU1200 significantly increased its output of viable offspring when starved as young adults (L4+12h and L4+24h) (Fig. 4A). KCNL-1 V530L may thus provide an immediate reproductive advantage depending on the environmental context. In addition, the JU1200 ARL_KCNL-1 V530L_ (but not JU751_WT_) displayed reduced embryonic lethality (Figs. 4B and S12B), indicating that internal hatching may shield embryos from external environmental insults, such as the imposed starvation stress. Environmental heterogeneity can therefore render the evolutionary transition from plastic to constitutive matricidal hatching via KCNL-1 V530L potentially beneficial. By competing JU1200_WT_ and JU1200 ARL_KCNL-1 V530L_ against a GFP-tester strain, we see that the V530L variant performed significantly worse in a constant food environment when the variant is introduced at frequencies of either 5% (Fig. S13A) or 50% (Fig. S13B).

**Figure 4.**
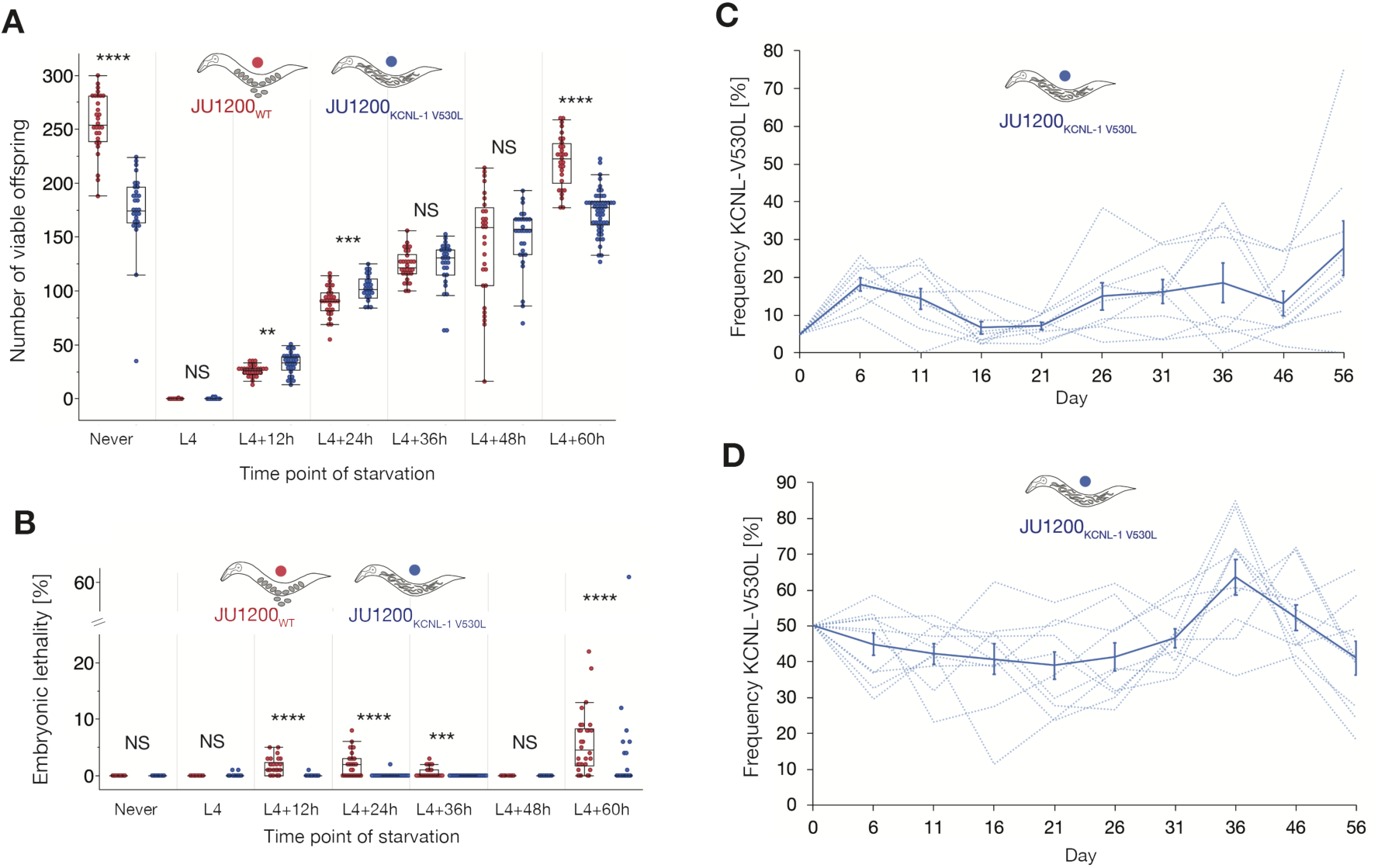
Fitness consequences KCNL-1 V530L in variable environments. (**A**) Lifetime production of viable offspring in selfing hermaphrodites in response to starvation encountered at varying maternal age: JU1200_WT_ versus JU1200 ARL_KCNL-1 V530L_. Age-synchronized populations were transferred from food (solid) to starvation (liquid) environment at different time points of development (N=24-53/strain/time point). Offspring number reflects combined viable larval offspring produced in nutrient-rich and starvation environments. (ANOVA separately performed for each time point: NS, not significant, \**P* < 0.05, \*\**P* < 0.01, \*\*\**P* < 0.001, \*\*\*\**P* < 0.0001). (**B**) Embryonic lethality in response to starvation encountered at varying maternal age: JU1200_WT_ versus JU1200 ARL_KCNL-1 V530L_ (from same experiment as in Fig. 4A, N=24-53/strain/time point). No embryonic lethality was observed in food conditions. (ANOVA separately performed for each time point: NS, not significant, \**P* < 0.05, \*\**P* < 0.01, \*\*\**P* < 0.001, \*\*\*\**P* < 0.0001). (**C**) Invasive capacity of JU1200 ARL_KCNL-1(V530L)_ into a JU1200_WT_ population at a starting frequency of 5% across ∼15 generations (N=9 replicates). Frequencies of the V530L allele are indicated by dashed lines (replicates), and the bold line indicates mean frequencies and standard error. (**D**) Direct competition of JU1200_WT_ versus JU1200 ARL_KCNL-1 V530L_ at an initial 1:1 ratio across ∼15 generations (N=10 replicates). Frequencies of the V530L allele are indicated by dashed lines (replicates), and the bold line indicates mean frequencies and standard error.

Once the derived JU751 allele escaped loss by genetic drift (Chelo et al., 2013), it could have been favoured by selection. To address this hypothesis, we tested whether the V530L allele (JU1200 ARL_KCNL-1 V530L_) can invade a JU1200_WT_ population when introduced at starting frequency of 5% (so that it is not lost by genetic drift) in a multiple-generation regime of fluctuating starvation stress (Fig. 4C). Across ∼15 generations, we found that the V530L allele was maintained at a mean frequency of 10-20%, although it was not detectibly beneficial across the experiment (χ^2^-test, 0.0139 ± 0.009, P=0.166), and only one of the nine replicate populations went extinct (Fig. 4C). We further quantified the effects of fluctuating starvation stress by direct competition of JU1200_WT_ versus JU1200 ARL_KCNL-1 V530L_ at a starting ratio of 1:1 across ∼15 generations (Fig. 4D), mimicking a situation where the JU751 allele was already established in the ancestral population. We found that throughout the competition assays, the KCNL-1 V530L allele was maintained at a mean frequency of 40-60% (Fig. 4D), again with little evidence that it was beneficial across the experiments (χ^2^-test, 0.006 ± 0.004, P=0.167). These results suggest that the KCNL-1 V530L allele is either neutral or that it is maintained by balancing selection in environments that vary in nutrient availability or other stress factors, similar to known environmental conditions of the *C. elegans* natural habitat that exhibit marked boom-and-bust population dynamics (Frézal and Félix, 2015).

The occurrence of constitutive *C. elegans* matricidal hatching in natural populations appears paradoxical given its highly deleterious effects on fecundity and survival. However, our data suggests that large-effect variants, such as KCNL-1 V530L, can be maintained, perhaps by selection, and that such extreme phenotypes uncovered in artificial conditions may be partly or completely masked in natural populations due to environmental heterogeneity (Taylor et al., 2019). We also note that the KCNL-1 V530L variant does not simply disrupt or abolish egg-laying as isolates are still capable of laying eggs albeit at a much-reduced rate. Hence, this variant strongly dampens environmental sensitivity of the egg-laying circuit, so that food signals usually triggering egg-laying cannot translate into sufficient or well-coordinated vulval muscle activation, which then leads to matricidal hatching.

Constitutive matricidal hatching in *C. elegans* caused by KCNL-1 V530L provides a rare example illustrating how a specific single-nucleotide DNA change underlies the evolution of a fixed, constitutively-expressed phenotype, ancestrally only induced in response to the environment. This observation is reminiscent of the phenomenon of genetic assimilation, *i.e.* the evolutionary process whereby an originally environmentally induced phenotype becomes genetically fixed and environmentally insensitive (Waddington, 1942). Yet, as we show, the mechanisms underlying phenotype fixation in the case of matricidal hatching reflect the acquisition of a novel single mutation, contrasting Waddington’s perceived idea of genetic assimilation by means of canalizing selection on cryptic genetic variation (Loison, 2019; Waddington, 1942, 1959). Importantly, however, recent experimental evidence suggests that at least some of Waddington’s *Drosophila* experiments involved stress-induced *de novo* mutations, which led to “pseudoassimilation” of selected phenocopies (Fanti et al., 2017). These and other discrepancies surrounding the role of phenotypic plasticity in evolution emphasize the need for more empirical work aimed at dissecting the molecular basis of evolutionary transitions between environmental and genetic control of phenotype expression.

## MATERIALS AND METHODS

### Strains and culture conditions

*C. elegans* strains were maintained on 1.7% or 2.5% agar NGM (Nematode Growth Medium) plates seeded with the *E. coli* strain OP50 at 15°C or 20°C (Brenner, 1974; Stiernagle, 2006). Experiments were performed at 20°C unless noted otherwise. The following strains were generated in this study: NIC1627 *kcnl-1(cgb1005[L530V*]) (allelic replacement line in JU751), NIC1642 *kcnl-1(cgb1002[V530L])* (allelic replacement line in JU1200), JIP1739 *kcnl-1(bln488[kcnl-1::d10])*, JIP1777 *kcnl-1(bln508[kcnl-1::SL2::wrmScarlet])*,JIP1803 *blnEx211[pSEM152 Pkcnl-1c::kcnl-1c(V530L)::SL2::GFP]*. For a complete list of strains used, including recombinant inbred lines and near-isogenic lines, see Table S1. Strains were thawed to allow for a maximum of four to six generation before phenotypic quantification to avoid accumulation of mutations. Strains were bleached in the second generation after thawing and kept in *ad libitum* food conditions on NGM plates until phenotypic observations.

### Experimental environments

We defined the standard culture condition (NGM agar plate seeded with the *E. coli* strain OP50) as a benign, favourable environment, in which *C. elegans* only rarely displays matricidal hatching, and if so, mostly late in life due to defects in the vulva(Pickett and Kornfeld, 2013). To assess matricidal hatching under starvation, we used liquid media (M9 or S-Basal) (Stiernagle, 2006) without adding bacterial food, and gentle agitation using a shaker. Starvation was performed in liquid cultures as *C. elegans* will leave agar plates without food, thus preventing phenotypic observations. Importantly, matricidal hatching is increased in liquid conditions even in the presence of ample food (Fig. S1A), likely due to increased egg retention caused by changes in oxygen levels and osmolarity, i.e. conditions known to affect egg retention (Schafer, 2006; Trent, 1982; Zhang et al., 2008). Therefore, the effects of starvation liquid culture on matricidal hatching observed here include other environmental factors in addition to starvation (e.g. mild hypoxia, increased osmolarity). This experimental environment can thus be considered an adverse, unfavourable environment due to multiple stressors. Liquid starvation treatment was imposed using 24-well plates (0.5 mL) or 96-well plates (0.1 mL) or by adding liquid medium directly onto starved NGM agar plates.

### Age-synchronization and developmental staging

Mixed-age hermaphrodite stock cultures kept at 20°C were hypochlorite-treated to obtain age-synchronized, arrested L1 populations. Hermaphrodites were then picked at the mid-L4 stage based on the morphology of the vulval invagination (Mok et al., 2015).

### Quantification of offspring number *in utero* and matricidal hatching

Offspring number (eggs and larvae) *in utero* was measured in randomly picked age-synchronized hermaphrodites, mounted directly on slides and gently squashed with a coverslip. The number of offspring *in utero* per individual was then counted using a 20x DIC microscope objective. An individual was considered to display matricidal (internal) hatching when one or more L1 larvae was visible in the uterus.

### Determination of embryonic stages

Embryonic stages were determined as described earlier(Sulston et al., 1983). The following ranges of embryonic stages were distinguished: early embryo (< 44-cell stage), intermediate embryo (> 44-cell stage < Pretzel (3-fold) stage), late embryo (> Pretzel (3-fold) stage) (Fig. S1B).

### Egg-laying behavioural assays

Egg-laying behavioural assays (Fig. S4) were performed by picking individual animals (L4+24h) to seeded NGM plates, using a modified, previously published protocol(Waggoner et al., 1998). Using a dissecting microscope, plates were scored every 2 minutes for the presence and number of eggs laid for a total duration of three hours. Eggs were immediately removed after egg-laying. Therefore, we only determined time intervals between active and inactive states of egg-laying and not intervals between individual eggs laid within active states, i.e. intra-cluster intervals (Waggoner et al., 1998).

### Quantification of hermaphrodite self-sperm number

We quantified the number of self-sperm in synchronized young adult hermaphrodites, i.e. adults containing one to six embryos *in utero* (Poullet et al., 2016). All animals had been isolated at the L4 stage to prevent mating with males. Individuals were collected in S-basal solution (5.85g/L NaCl, 1g/L K2HPO4, 6g/L KH2PO4, 5mg/L cholesterol) and fixed in cold methanol (−20°C) for at least 30 minutes, washed three times with PBTw (PBS: phosphate-buffered saline, 137 mM NaCl, 2.7 mM KCl, 10 mM Na2HPO4, 2 mM KH2PO4, pH 7.4 containing 0.1% Tween 20) and squashed on a glass slide with Vectashield mounting medium containing 4,6-diamidino-2-phenylindole (DAPI). Imaging of the anterior spermatheca was performed using an Olympus BX61 microscope with a CoolSnap HQ2 camera. Images were taken at 60X magnification as Z-sections (1 µm) covering the entire gonad. Sperm number was counted by identifying condensed sperm nuclei on each focal plane using the Fiji plugin Cell Counter. When primary spermatocytes were still present, they were counted as four sperm. We assessed sperm number in a single gonad arm of each individual. Sperm number determined from one gonad arm was multiplied by two to infer the total number of sperm produced.

### Quantification of offspring production and embryonic lethality

Lifetime production of viable offspring was analysed by isolating mid-L4 hermaphrodites on individual NGM plates and transferring them daily to fresh NGM plates until egg-laying ceased. The number of live (viable) larvae and unhatched eggs (embryonic lethality) was counted 24-36 hours after each transfer. When mothers died due to matricidal hatching, trapped larvae were allowed to exit the dead mother before being counted. Alternatively, mothers were squashed between slide and coverslip for counting of offspring in the uterus. In liquid culture, offspring production was measured by counting larvae and eggs in wells; eggs that failed to hatch after 24-36 hours were considered to be unviable (embryonic lethality). To count offspring *in utero*, animals in liquid cultures were transferred to a microscope slide using an eyelash. All strains were scored in parallel.

### Lifespan analysis

Young adult hermaphrodites (L4+24h) were transferred to NGM plates seeded with fresh *E. coli* OP50 bacteria. Individuals were transferred daily onto fresh plates until the end of egg laying and every three days after that until the experiment was completed. Animals were scored as dead if they failed to respond to a tip on the head and tail with a platinum wire. For each temperature regime (15°C and 20°C), the two strains were scored in parallel.

### Embryo size

Laid embryos of age-synchronized hermaphrodites (at L4+24h and L4+48h) were collected using M9 mounted on 4% agar pads on glass slides and images were obtained with ImageJ software using a 40x DIC microscope objective. The length and width of each egg shell was measured to calculate embryo volume with the ellipsoid volume function: 4/3 x π x (length/2) x (width/2)^2^.

### Body size

Body size of age-synchronized mid-L4 hermaphrodites was measured using digital images take via a 20x DIC objective. As an estimate of body size, we measured the perimeter of animals by manually tracing the outline of the body in the Fiji software.

### Serotonin assays

Based on previously described protocols (Horvitz et al., 1982; Trent et al., 1983; Waggoner et al., 2000), young adults (mid L4+24h) were transferred to individual wells of a 96-well microtiter plate containing 100 µL of M9 buffer (without bacterial food) containing 10 mg/mL serotonin (5-hydroxytryptamine creatinine sulfate complex, Sigma). Egg-laying rates were measured by counting the number of eggs laid after three hours in the presence or absence of serotonin at 20°C (Fig. 1H). All strains were scored in parallel.

### Defecation assays

Defecation cycle length of singled hermaphrodites was measured at mid-L4+24h (Fig. S9C). Cycle length was determined as the time interval between the first muscular contraction of a defecation event and the first muscular contraction of the next (Wong et al., 1995). For each individual, five defecation cycles were measured.

### Generation of F2 Recombinant Inbred Lines (RILs)

We generated 144 F2 Recombinant Inbred Lines based on a reciprocal parental cross between wild isolates JU751 (Le Perreux-Sur-Marne, France, isolated from garden compost in 2005) and JU1200 (Scotland, UK, isolated from garden compost in 2007) (Fig. S5A). Prior to crossing of parental isolates, hermaphrodites were depleted of their self-sperm to obtain only cross-progeny in the F1 generation. RILs were derived from 36 randomly picked F2 individuals among F1 cross-progeny obtained per cross direction and parental origin (fig. S5A). The total of 144 F2 individuals were then inbred for 12 generations through selfing by randomly picking a single animal in each generation (Fig. S5A).

### Genotyping of RILs

DNA from the two parental strains (JU1200, JU751), the two F1 reciprocals and 144 RILs was prepared using the Qiagen Blood and Tissue kit soon after derivation or after thawing from frozen stocks and expansion to at least 10^3^ L1 individuals. Genotyping of 164 bi-allelic single-nucleotide polymorphisms (SNPs), uniformly distributed across the six chromosomes according to published genetic distances (Rockman and Kruglyak, 2009) and parental identities(Andersen et al., 2012), was done with the iPlex Sequenom MALDI-TOF platform (Bradić et al., 2011). For each sample, a multi-plex PCR targeting between 30 and 35 SNP regions, was followed by a SNP allele-specific extension and genotyping by mass spectrometry. A list of the SNPs targeted and the oligonucleotides employed for Sequenom analysis can be found in Table S2. A total of 164 SNP markers were genotyped of which 146 SNPs were retained for QTL mapping after quality control. SNP quality control first involved retaining those where either the two parental strains had a different allele or one of the two reciprocal F1s was heterozygous. The recombinant genotypes of RILs are listed in the supplementary file Data S1.

### Phenotyping of RILs

Across multiple experimental blocks, we scored matricidal (internal) hatching at 15°C. For each RIL (N=144), 3-4 replicates with each 30 mid-L4 were established and after 24h, 48h and 72h, the proportion of individuals containing one or more internally hatched larva was scored (Fig. 2B and Fig. S5B). Parental isolates were scored in parallel in each block. For QTL mapping, we used the cumulative proportion of animals displaying matricidal hatching on day 3.

### QTL analysis

QTL mapping was performed using the rQTL package in R (Broman et al., 2003). The cross data was converted into recombinant inbred lines using the convert2riself function. After visual inspection of the genetic map and recombination fractions as well considering the values from the top.errorlod and droponemarker functions, we removed four markers due to them creating substantial expansions of the genetic map and having anomalous recombination patterns (chromosome 3: III_8052645[C/A], chromosome 5: V_7531256[A/G], V_1297644[C/G], V_3936556[A/G]).

The 5% significance threshold for the QTL mapping was calculated using 1,000 permutations. The confidence interval for the QTL on chromosome 5 was calculated using the bayesint function in rQTL using a probability of 0.95.

### Construction of near-isogenic lines (NILs) and fine-mapping

To validate the effect of the uncovered QTL in *C. elegans* matricidal hatching, we derived near-isogenic lines from four RILs (NIC613, NIC632, NIC670, NIC707) displaying constitutive matricidal hatching and different break-points within the QTL region (Fig. S7B). Corresponding NILs (NIC1566, NIC1570, NIC1573, NIC1577) were established by eight rounds of backcrossing to the parental JU1200 isolate selecting for individuals with matricidal hatching/markers. NILs were then phenotyped and genotyped (Fig. S7).

### Molecular Biology

pSEM148 was built by assembling the following DNA fragments using Gibson assembly: (1) LHR (left homology arm), (2) SL2::wrmScarlet cassette, (3) RHR (right homology arm). pSEM152 was built similarly by joining (1) the 2.8kb sequence upstream of the *kcnl-1c* start codon, (2) the *kcnl-1c* locus including the V530L mutation, (3) and SL2::GFP cassette. pCFJ55 plasmids were generated by three-fragment Gateway assembly from: (1) [4-1] Pkcnl-1 isoform C (2.9 kb), (2) [1-2] kcnl-1 isoform C [A = wt, B = A443V], and 3 [2-3] GFP unc-54 UTR. See Table S2 for detailed reagents.

### *C. elegans* CRISPR-*Cas9* gene editing and transgenesis

Gene editing was performed according to previously published protocols(Dickinson and Goldstein, 2016; El Mouridi et al., 2017) (see Table S2 for detailed reagents). In brief, *bln488* was built by inserting the *d10* protospacer sequence at the C-terminus of the wild-type *kcnl-1* locus using the crRNA *kcnl-1_Cter* and the single-strand oligonucleotide repair template (ssODN-RT) oSEM474. *bln508* was generated by using the *bln488 d10* locus as a starting point and engineering the *SL2::wrmScarlet* cassette using the pSEM148 repair template plasmid.

The *blnEx211([Pkcnl-1c::kcnl-1c(V530L)::SL2::GFP])* transgene was generated by injecting the pSEM152 plasmid and the pCFJ90 (Pmyo-2::mCherry) co-injection marker into the wild-type N2 strain.

The KCNL-1 L530 residue found in the wild isolate JU751 was mutated to V530 (*i.e. cg1005*) using the crRNA *kcnl-1_cg1005* and ssODN-RT *kcnl-1_L530*, resulting in NIC1627. In brief, JU751 young adult worms were injected with Cas9 protein, *kcnl-1* and *dpy-10* crRNA and ssODN-RT. This injection mix was prepared in the following way: (i) combine 3.9 µl of Cas9 protein (Dharmacon, 20 µM), 1.25 µL tracrRNA (Dharmacon, 0.175 mM), 0.2 µL *dpy-10* crRNA (0.3 mM), 0.25 µL *kcnl-1* crRNA (*kcnl-1_cg1005*, 0.6 mM); (ii) Incubate 15 min at 37°C; (iii) add 0.2 µL *dpy*-10 ssODN-RT (AF-ZF-827mod, 0.011 mM) and 0.2 µL *kcnl-1* ssODN-RT (*kcnl-1_L530*). F1 worms were isolated from P0 plates containing rollers, and allowed to produce offspring for 2 days for subsequent use (F2 progeny). The original F1 worms were then individually lysed and screened by PCR using three oligonucleotides: kcnl-1_FWD, 5’ of the SNP; kcnl-1_REV, 3’ of the SNP; kcnl-1_SNP that carries the modified SNP.

The KCNL-1 V530 residue found in the wild isolate JU1200 was mutated to L530 (*i.e. cg1002*) using the crRNA *kcnl-1_cg1002* and ssODN-RT *kcnl-1_V530*, resulting in NIC1642. *cg1002* animals were detected in the F1 generation based on their egg retention phenotype, and verified in the F2 by Sanger sequencing.

### Focus of action of *kcnl-1*

The focus of action of *kcnl-1* was determined by correlating the presence/absence of the *blnEx211* transgene in vulval muscle and VC neurons with the egg-laying phenotype (wild-type or bag of worms) of individual worms raised at 25°C, over the course of 48 hours after the final larval moult.

### RNA interference

We used an RNAi clone targeting the *kcnl-1* gene (clone B0399.1) from the Orfeome RNAi library(Rual et al., 2004) to test if RNAi by feeding (Kamath et al., 2003) could suppress the egg-laying defect of the JT6428 (*exp-3*(*n2372*)) strain. As a negative control, we cloned a 519 bp fragment of eGFP into the pDEST-pL4440 RNAi vector that contains two concordant T7 promoters. We transferred five young adult hermaphrodites to plates with RNAi bacteria and removed all adults after three hours. Three days later we counted the number of adult F1 progeny on the plate. Experiments were performed at room temperature in duplicate on three to five plates.

### Overexpression of mutant KCNL-1 protein

To test the causal role of the *exp-3(n2372)* (A443V) mutation identified in the *kcnl-1* gene, we generated wild-type or mutant GFP tagged cDNAs of the C isoform of *kcnl-1*. We generated expression constructs using a 2876 bp promoter for the C isoform of *kcnl-1* and a 3’ UTR from *unc-54* by three-fragment Gateway cloning (Thermo Fisher, USA). See Table S2 for details. We generated extra-chromosomal array lines by injection into *lin-15*(*765ts*) animals using the Plin-15EK rescue plasmid (Clark et al., 1994) at 150 ng/ul and the *kncl-1* expression constructs (wildtype or mutant) at 10 ng/ul. We tested egg-laying in three independent lines (EG4318, EG4319, EG4320, EG4321) carrying wild-type or mutant *kcnl-1* by allowing five young adult transgenic animals to lay eggs for 3 hours in triplicate and counting the number of adult F1 progeny three days later.

### Reproduction in response to starvation encountered at varying maternal age

Age-synchronized mid-L4 hermaphrodites from either JU1200_WT_, JU1200 ARL_KCNL-1 V530L_ JU751_WT_ and JU751 ARL_KCNL-1 L530V_ strain were isolated and cultured on NGM plates seeded with *E. coli* OP50. Except for the “never” treatment group, which was not exposed to starvation, individuals were transferred to M9 buffer in 12-well plates with gentle shaking at 20° (1 animal per well, 1ml M9 per well, N∼30 individuals per treatment group) for all other treatment groups (starvation at 0 h, 12 h, 24 h, 36 h, 48 h, 60 h). The liquid starvation treatment was applied for 48 hours, during which virtually all animals, except the ones from treatment group “0h” had died within the first 24 hours. For the treatment group “0 h”, individuals were directly transferred to individual wells using an eyelash. For the treatment groups “12 h”, “24 h”, individuals were transferred to individual wells 12 h or 24h after isolation and the NGM plates of origin were conserved for subsequent counting of the offspring laid by each individual before its transfer to liquid. For the treatment groups “36 h”, “48 h” and “60 h”, individuals were passaged each day to fresh NGM plates before transfer to the starvation treatment. Plates were conserved for subsequent counting of the offspring prior to starvation. The treatment group “never” was maintained on NGM plates until the cessation of reproduction, i.e. L4 +96 h) and all individuals were daily transferred to fresh NGM plates. Viable larval offspring and unhatched (dead) embryos produced during the 48h starvation treatment were counted using a dissecting microscope. Offspring trapped in dead adult bodies were transferred to slides using an eyelash for microscopy observation using a 20x DIC objective. Larval offspring unhatched (dead) embryos produced on NGM plates prior to starvation were counted 24-36h after transfer of individuals. Total numbers of viable offspring shown in Figs. 4A and S12A include the lifetime production of viable larval offspring produced on NGM plates and liquid starvation. Embryonic lethality in liquid starvation (Figs. 4B and S12B) was inferred from the number of unhatched eggs present in wells after 48 h of the treatment. No embryonic lethality was observed for any strain on NGM plates.

### Competition and invasion experiments

#### *Competition with GFP-tester strain in ad libitum food conditions* (Fig. S13)

To evaluate long-term fitness consequences of KCNL-1 V530L, we first performed competition experiments in a favourable, constant *ad libitum* food environment across multiple generations. We ran competition experiments for both JU1200_WT_ and JU1200 ARL_KCNL-1_ V530L against a GFP-tester strain (*myo-2::gfp*; strain PD4790, expressing green fluorescent protein (GFP) in the pharynx) with a starting frequency of either 5% (invasion; fig. S13A) or 50% (competition; Fig. S13B). To establish experimental populations, age-synchronized arrested L1 larvae of different strains were mixed to obtain a total number of 1000 larvae per NGM plate. For the invasion experiments with a starting ratio of 5:95, we established (a) mixed populations containing 50 (L1 larvae of JU1200_WT_ and 950 L1 larvae of *myo-2::gfp* tester strain (N=10 replicates) and (b) mixed populations containing 50 L1 larvae of JU1200 ARL_KCNL-1 V530L_ and 950 L1 larvae of *myo-2::gfp* tester strain (N=10 replicates) (Fig. S13A). For the competition experiments with a starting ratio of 1:1, we established (a) mixed populations containing 500 L1 larvae of JU1200_WT_ and 500 L1 larvae of *myo-2::gfp* tester strain (N=10 replicates) and (b) mixed populations containing 500 L1 larvae of JU1200 ARL_KCNL-1 V530L_ and 500 L1 larvae of *myo-2::gfp* tester strain (N=10 replicates) (Fig. S13B). Populations were maintained for 20 days by daily chunking of agar carrying nematodes (approximately one quarter of the agar) to fresh NGM plates. Every four days, genotype frequencies were inferred by quantifying the fraction of GFP-positive individuals among a subpopulation of ∼200-300 individuals per replicate, using a fluorescence dissecting microscope.

#### *Direct competition in fluctuating starvation environments* (Figs. 4C and 4D)

We performed direct competition experiments of JU1200 ARL_KCNL-1 V530L_ versus JU1200_WT_, thus assessing the effects of the single amino acid change (KCNL-1 V530L) in an identical genetic background (JU1200). After initial expansion of the populations, the culturing regime alternated between 72 hours on NGM plates (with food) and 48 hours in liquid (without food). To establish invasion experiments at a 5:95 ratio (Fig. 4C), we mixed 3 mid-L4 JU1200 ARL_KCNL-1 V530L_ and 57 mid-L4 JU1200_WT_ individuals per standard NGM agar plate seeded with *E. coli* OP50 (N=9 replicates). We established competition experiments at 1:1 ratio (Fig. 4D) by mixing 10 mid-L4 JU1200 ARL_KCNL-1 V530L_ and 10 mid-L4 JU1200_WT_ individuals per standard NGM agar plate seeded with *E. coli* OP50 (N=10 replicates). After 24h, for both invasion and competition experiments, all replicate populations were transferred to the starvation treatment (5 mL M9 liquid culture without bacterial food source in 15 mL Falcon tubes, with gentle shaking) for 48h. At this time, we added 0.5 mL of the starved liquid culture to fresh NGM plates seeded with *E. coli* OP50. Populations were then allowed to grow for 72 hours. The food-starvation alternation was then continued for a total of 10 cycles across 56 days. Allele frequencies were determined by phenotyping a random subset (N=100) of hermaphrodites (at L4+48h) of each replicate population (days 16, 21, 26, 31, 36, 46, 56). As validated by PCR genotyping, individuals with >25 embryos *in utero* could be reliably scored as JU1200 ARL_KCNL-1 V530L_ whereas JU1200_WT_ individuals contained < 20 embryos *in utero.* Exceptional cases (N=4) where individuals had >20<25 embryos *in utero* were excluded from analysis. In addition, a random subset of individuals (N=100) from each replicate population was genotyped at the KCNL-1 locus using PCR primers discriminating between V530 and L530 on days 6 and 46 of the experiment.

#### Estimation of allele frequency dynamics

*kcnl-1* allele frequency dynamics were analysed using a generalized linear mixed model in R(R Development Core Team 3.0.1., 2013), using the *lme4* package(Bates et al., 2015) to estimate the relative fitness of JU751 allele (Chelo et al., 2013). The two competition experiments were analysed separately as they were done at different starting allele frequencies. *kcnl-1* allele counts were modelled as a function of fixed replicate intercept and fixed day of competition while taking the slope of replicate as a random binomial variable. We ignored allele frequencies at set-up since they there is a positive correlation between replicate intercept and slope. To test for significance of day of competition – whose estimate is the relative fitness of the JU751 allele – a second model without day as a fixed factor was compared with the first model using a chi-squared test.

### Statistical analyses

Statistical tests were performed using R and JMP 14.0. Data for parametric tests were transformed where necessary to meet the assumptions of ANOVA procedures (homogeneity of variances and normal distributions of residuals). For *post hoc* comparisons, Tukey’s honestly significant difference (HSD) procedure was used. For data, where ANOVA assumptions could not be met, we used nonparametric tests (*e.g.* WilcoxonTest).

Box-plots: the median is shown by the horizontal line in the middle of the box, which indicates the 25th to 75th quantiles of the data. The 1.5 interquartile range is indicated by the vertical line.

## Supporting information

Data S1 (raw data)

## DATA AVAILABILITY

All raw data are provided in the supplementary file Data S1. The authors affirm that all data necessary for confirming the conclusions of the article are present within the article, figures, and tables.

## ACKNOWLEDGEMENTS

CB, CF, CG and PV acknowledge financial support by the Centre National de la Recherche Scientifique (CNRS), the Institut national de la santé et de la recherche médicale (Inserm), and the Université Côte d’Azur (UCA). HT and AP-Q were supported by the European Research Council (FP7/ 2007-2013/243285) and the Agence Nationale de la Recherche (ANR-14-ACHN-0032-01). JH was supported by the French Government (National Research Agency, ANR) through the “Investments for the Future” LABEX SIGNALIFE: program reference # ANR-11-LABX-0028-01. TB and SEM were supported by the European Research Council (337702-Kelegans). CFJ is supported by KAUST intramural funding. For helpful discussions and pointing out the curious phenotype of JU751, we would like to thank Marie-Anne Félix. For help with experimental work and data analysis, we thank Chami Kim, Bénédicte Billard, Sarah Fausett, Laure Mignerot, Nausicaa Poullet, Ghada Bouzouida, Nicolas Schwartz-Tamoglia, Nicolas Callemeyn-Torre, Romain Salle, João Costa and François Mallard. For advice and comments on previous versions of the manuscript, we thank Ehab Abouheif and Luke Noble. Strains and materials were provided by Marie-Anne Félix, the *C. elegans* Natural Diversity Resource (CeNDR) and the *Caenorhabditis* Genetics Center (CGC), which is funded by NIH Office of Research Infrastructure Programs (P40 OD010440).

## AUTHOR CONTRIBUTIONS

CB conceived and supervised the study, and wrote the manuscript. PV, CG, CF performed the majority of experiments, including construction of RILs, phenotyping, molecular biology and competition experiments. PV and CG edited the manuscript. AV contributed to construction of RILs and RIL genotyping. AP-Q and HT designed and conducted RIL genotyping. HT contributed to experimental design, analysed data and writing of the manuscript. JH analysed RIL genotype and phenotype data and performed the QTL analysis. TB and SEM performed *kncl-1* expression and mosaic analysis, and edited the manuscript. CFJ cloned *exp-3*, performed RNAi experiments, and edited the manuscript.

## Supporting Information

### SUPPLEMENTARY FIGURES

**Fig. S1.**
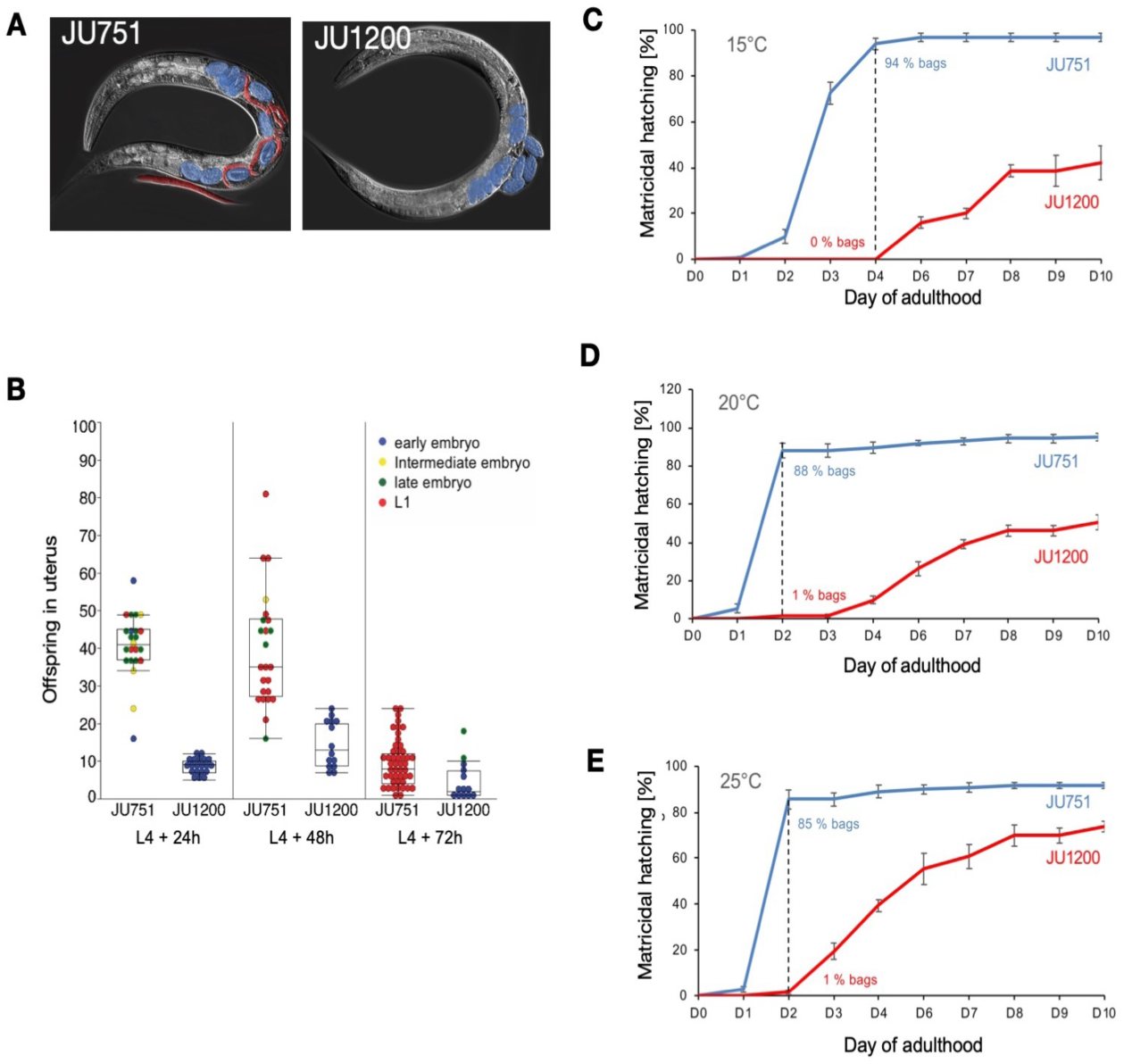
Constitutive matricidal hatching in JU751. (**A**) Left: Image of an adult JU751 hermaphrodite (mid-L4+48h) containing internally hatched larvae (red) and eggs (blue) with advanced embryonic stages, in food conditions. Right: Image of an adult JU1200 hermaphrodite (mid-L4+48h) with canonical *C. elegans* egg-laying behavior, containing eggs (blue) with early embryonic stages. (**B**) Number and developmental stages of offspring *in utero* of hermaphrodites in JU751 and JU1200 across the reproductive span of adulthood. At all time points, JU751 contains a much higher number of internally developing progeny at more advanced developmental stages. At the mid-L4+72h stage, all JU751 individuals contained at least one internally hatched larva. Each data point reflects the number of internal offspring developing within a single individual (N=14-47 individuals/strain/time point). Color indicates the developmental stage of the oldest progeny observed within the individual: early embryo: < 44-cell stage (blue); intermediate embryo: > 44-cell stage < Pretzel (3-fold) stage (yellow); late embryo: > Pretzel (3-fold) stage (green); L1 larva (red). Same dataset as in Fig. 1D-F. (**C-E**) Temporal dynamics of matricidal hatching in JU751 versus JU1200 at three different growth temperatures. Figures display cumulative percentages of individuals containing at least one internally hatched larva. The percentage of individuals with internal hatching (“bags”, *i.e.* containing at least one larva *in utero*) are indicated by a dotted line on selected days. The dataset of (**D**) is the same as shown in Fig. 1C.

**Fig. S2.**
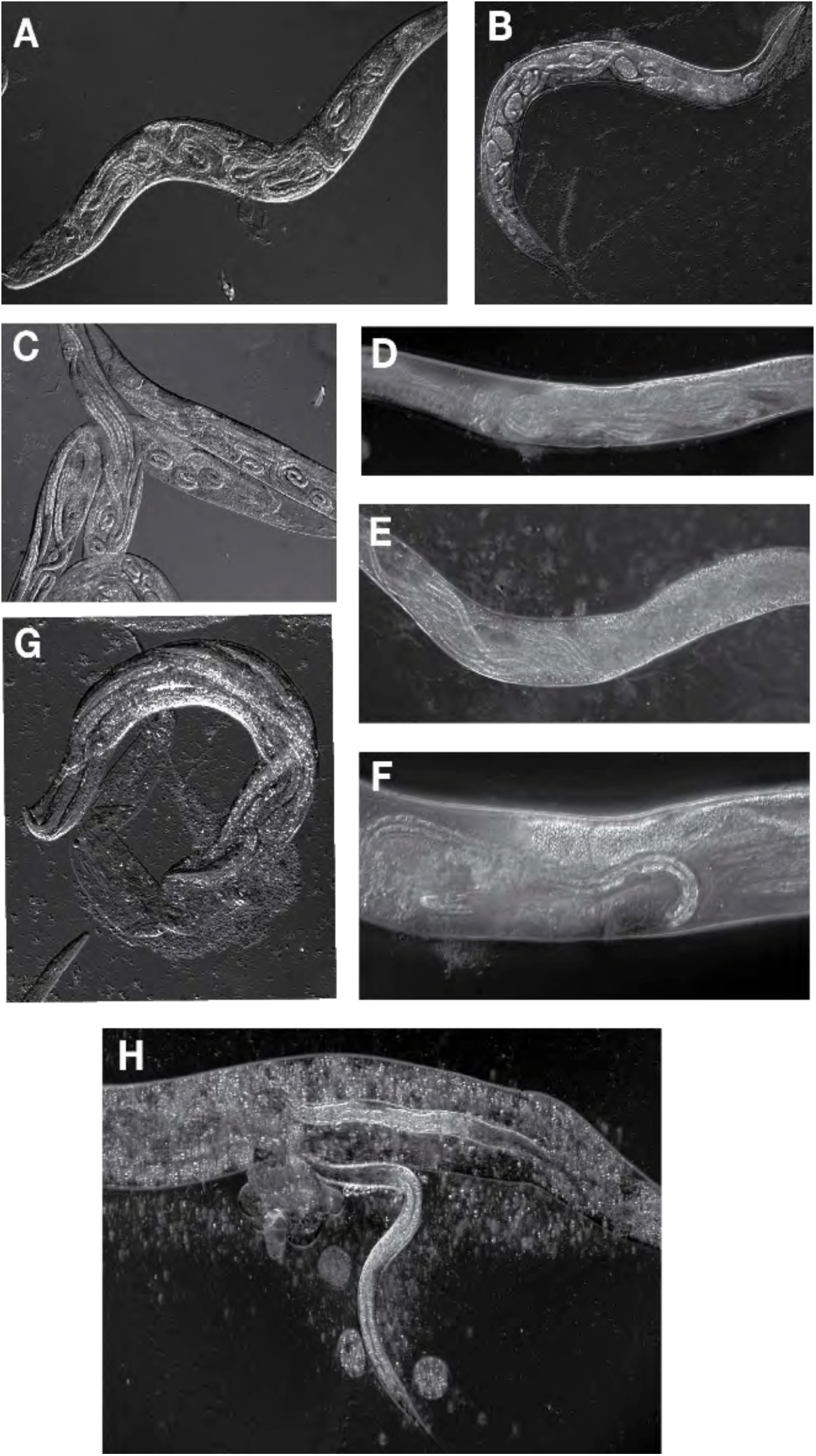
Early maternal death due to internal hatching in JU751. (**A**-**H**) DIC microscopy image of JU751 hermaphrodites on day three of adulthood (mid-L4+72h). (**A, B**) Live individuals containing larvae and embryos at advanced stage; internal organs and tissues, including germline, of mothers have been completely destroyed. (**C**-**G**) Dead individuals containing larvae and embryos at advanced stage. (**H**) Larva exiting the maternal cuticle (“bag”).

**Figure S3.**
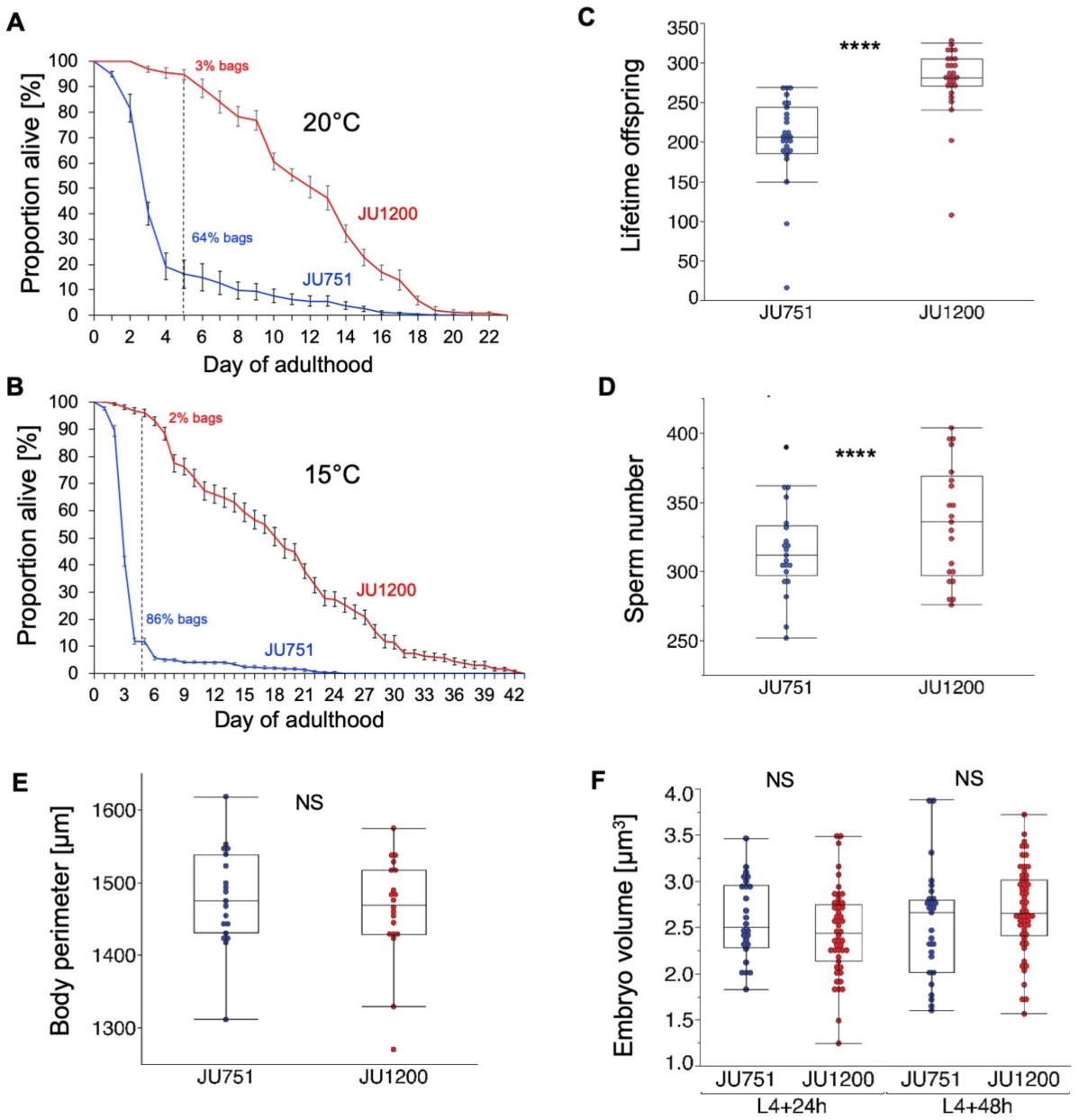
Life history differences between JU751 and JU1200. (**A**) Survival curves of JU751 and JU1200 hermaphrodites in *ad libitum* food conditions at 20° (same data as in Fig. 1G). The dotted line indicates the percentages of individuals with matricidal hatching (“bags”) on day five. (**B**) Survival curves of JU751 and JU1200 hermaphrodites in *ad libitum* food conditions at 15°. The dotted line indicates the percentages of individuals with matricidal hatching (“bags”) on day five. (**C**) Lifetime offspring production of selfing JU751 and JU1200 hermaphrodites (20°C). JU751 produces a strongly reduced number of offspring (198.2 ± 0.4) relative to JU1200 (275.7 ± 0.4) (ANOVA, F_1,53_=21.7, P<0.0001). (**D**) Sperm number of selfing JU751 and JU1200 hermaphrodites (20°C): no significant difference (ANOVA, F_1,40_=2.9, P=0.10). (**E**) Body size of JU751 and JU1200 hermaphrodites at the mid-L4 stage (20°C): no significant difference (ANOVA, F_1,37_=0.5, P=0.50).(**F**) Embryo size of JU751 and JU1200 hermaphrodites (20°C): no significant differences at L4+24h (ANOVA, F_1,73_=1.7, P=0.20) or L4+48h (ANOVA, F_1,79_=2.1, P=0.15).

**Fig. S4.**
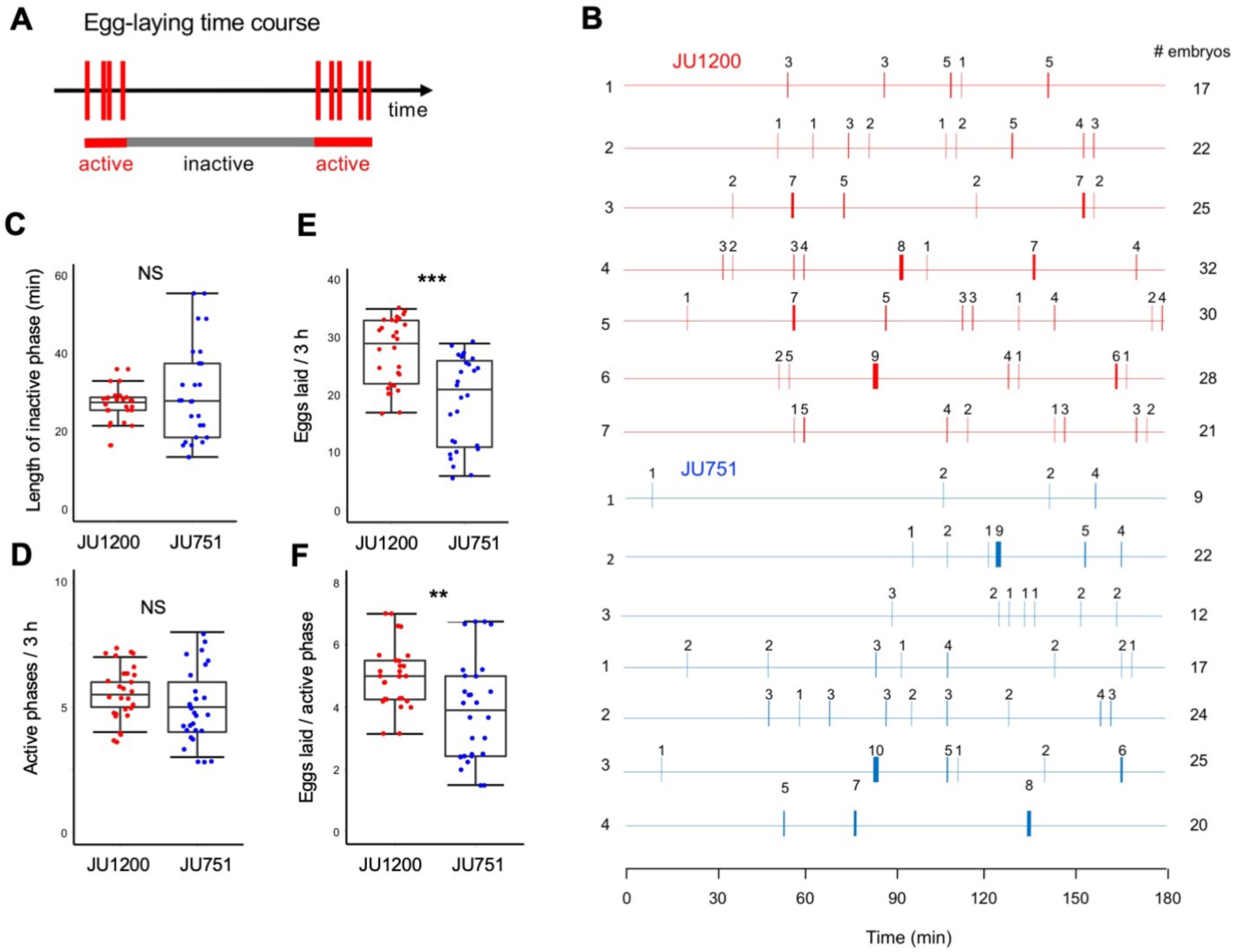
Egg-laying behavior in JU751 and JU1200. (**A**) Cartoon outlining the temporal patterns of *C. elegans* egg-laying behavior (Waggoner et al., 1998). *C. elegans* egg-laying is rhythmic, comprised of active egg-laying phases (∼2 minutes) interspersed by inactive phases of ∼20 minutes (gray), as measured in the reference strain N2 (Waggoner et al., 1998). Intervals within active states are called intra-cluster intervals and intervals between two active events are called inter-cluster intervals (Waggoner et al., 1998). Red bars indicate egg-laying events (number of eggs laid are shown above bars; thick bars indicate events with multiple eggs laid). In our experiments (**B-F**), we quantified the time of inter-cluster intervals and the number of eggs laid per active phase (but not the time of intra-cluster intervals). (**B**) Tracking of egg-laying phases in individual hermaphrodites (L4+24h) of JU751 and JU1200 across a 3-hour interval. Each line represents a single individual (N=7 per strain). A total of 28 individuals were scored per strain. (**C**) The variance (Levene’s test, F_1,54_=19.6, P<0.0001) but not the mean length (Wilcoxon Test, P=0.79) of inactive phases was significantly increased in JU751 relative to JU1200. **(D**) The mean number of active egg-laying phases was slightly lower in JU751 compared to JU1200 with marginal significance (Wilcoxon Test, P=0.06). (**E**) JU1200 laid a significantly higher total number of embryos compared to JU751 (Wilcoxon Test, P=0.0002). (**F**) JU1200 laid a significantly higher number of embryos per active phase compared to JU751 (Wilcoxon Test, P=0.0048).

**Fig. S5.**
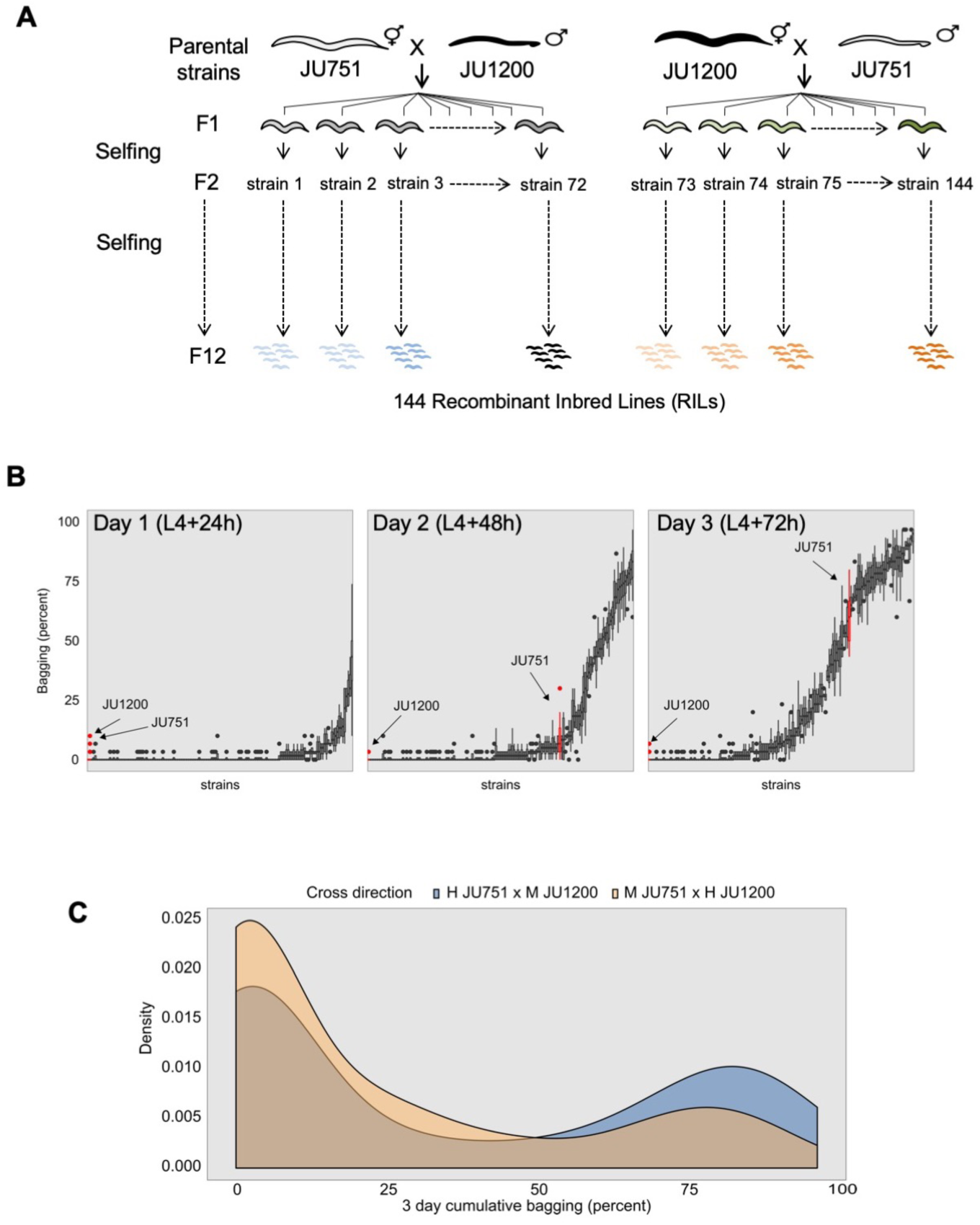
Generation and phenotyping of Recombinant Inbred Lines (RILs). (**A**) Detailed crossing scheme used to generate F2 RILs. (**B**) Phenotype distribution for all RILs (*x*-axis) shown as boxplots made up by the replicates of each RIL. Cumulative proportion of individuals displaying internal hatching (i.e. at least one L1 larva within mother) across the first three days of adulthood. Parental strains are colored in red. (**C**) Phenotype distribution for RILs sorted by their cross direction.

**Fig. S6.**
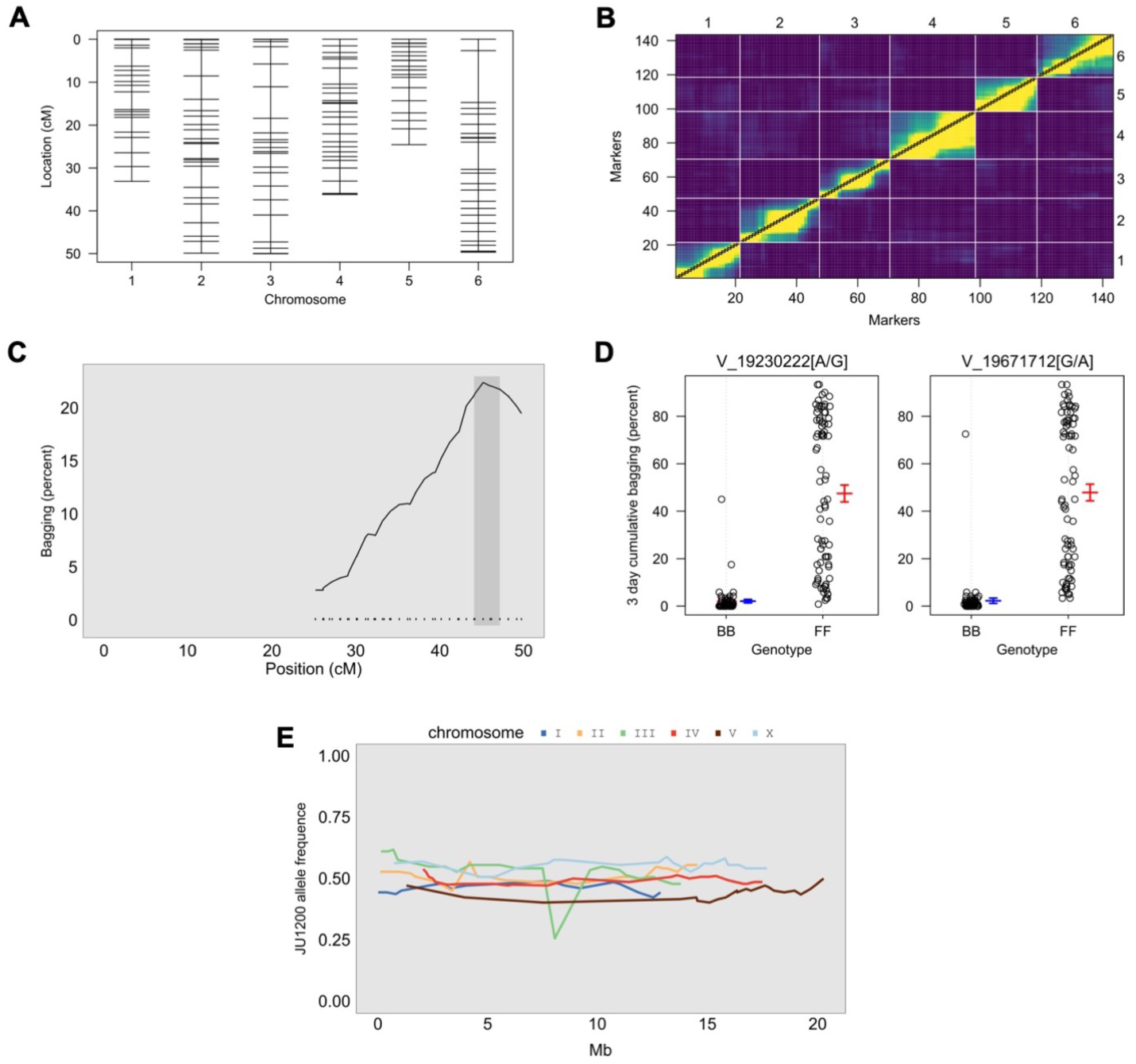
SNP genotyping, genetic map and QTL mapping. (**A**) Genetic markers cover the entire genome. (**B**) Heatmap of the recombination fractions (top) and LOD scores (bottom) for the genetic map, yellow corresponds to less recombination between markers while purple corresponds to more recombination between markers. LOD scores result from testing if the recombination fraction = 0.5, as per the est.rf() function in rQTL. (**C**) QTL peak on chromosome 5, the shaded area is the 95 percent confidence interval for the peak calculated with the bayesint function in rQTL. **(D**) Phenotypic distribution at the two markers underneath the QTL peak on chromosome 5. BB contains the strains that are homozygous for the JU1200 allele at that locus, FF contains the strains that are homozygous for the JU751 allele at that locus. (**E**) Allele frequencies for the JU1200 allele across all RILs, cluster around 50% as expected with a small decrease on chromosome 3 at around 8Mb.

**Fig. S7.**
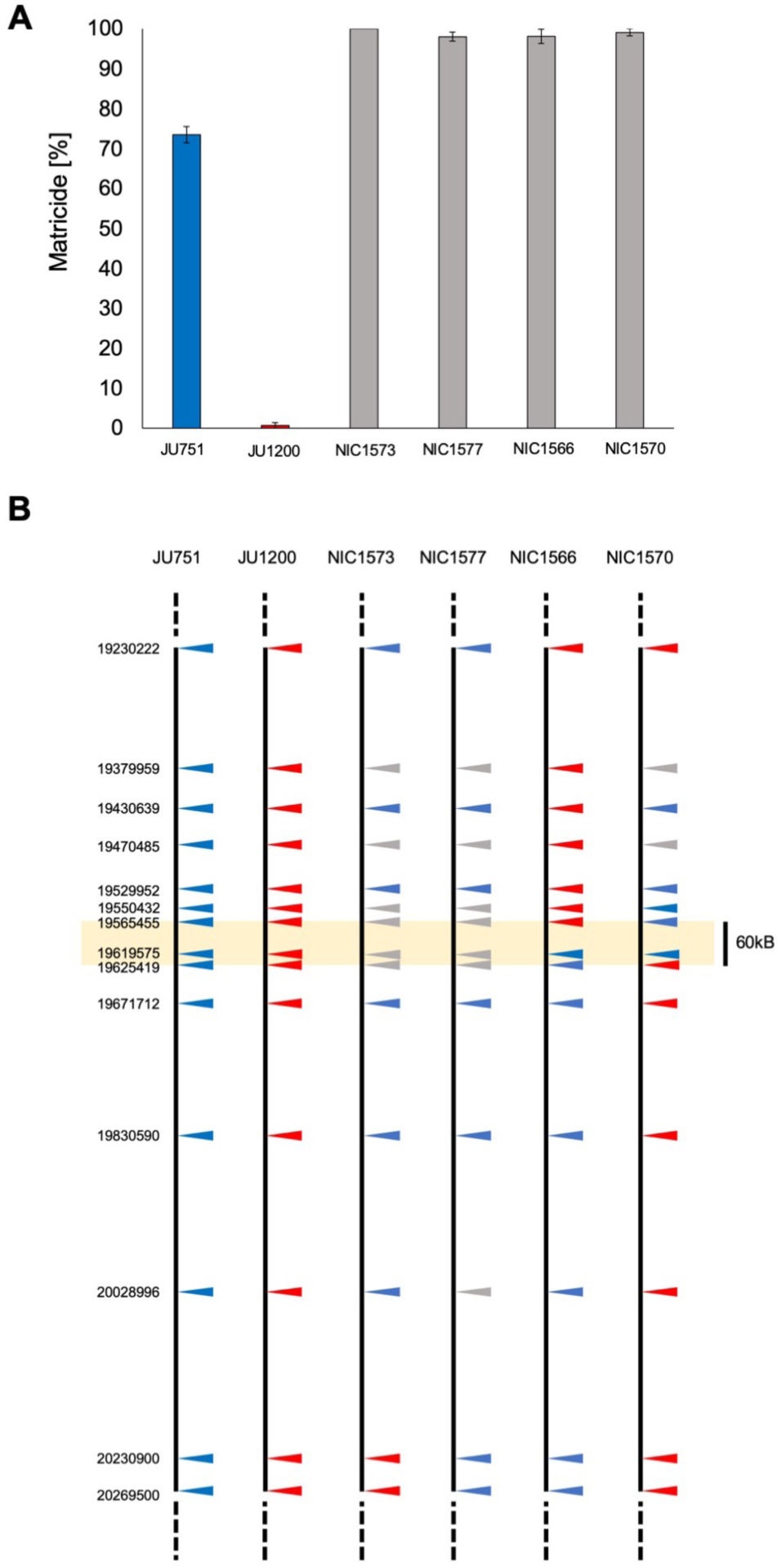
Phenotyping of near-isogenic lines and fine-mapping of QTL region. (**A**) Phenotyping of matricide in near-isogenic lines (NILs): values indicate cumulative percentage of individuals containing one or more internally hatched larvae after three days of adulthood (L4 + 72h) (N=3-5 replicates/strain, N=30 individuals/replicate). Parental isolates were scored in parallel. (**B**) Fine-mapping of the QTL region to a 60 kb region (yellow rectangle) based on NIL genotyping at SNP makers (triangles): red (JU1200 genotype), blue (JU751 genotype), gray (unknown genotype).

**Fig. S8.**
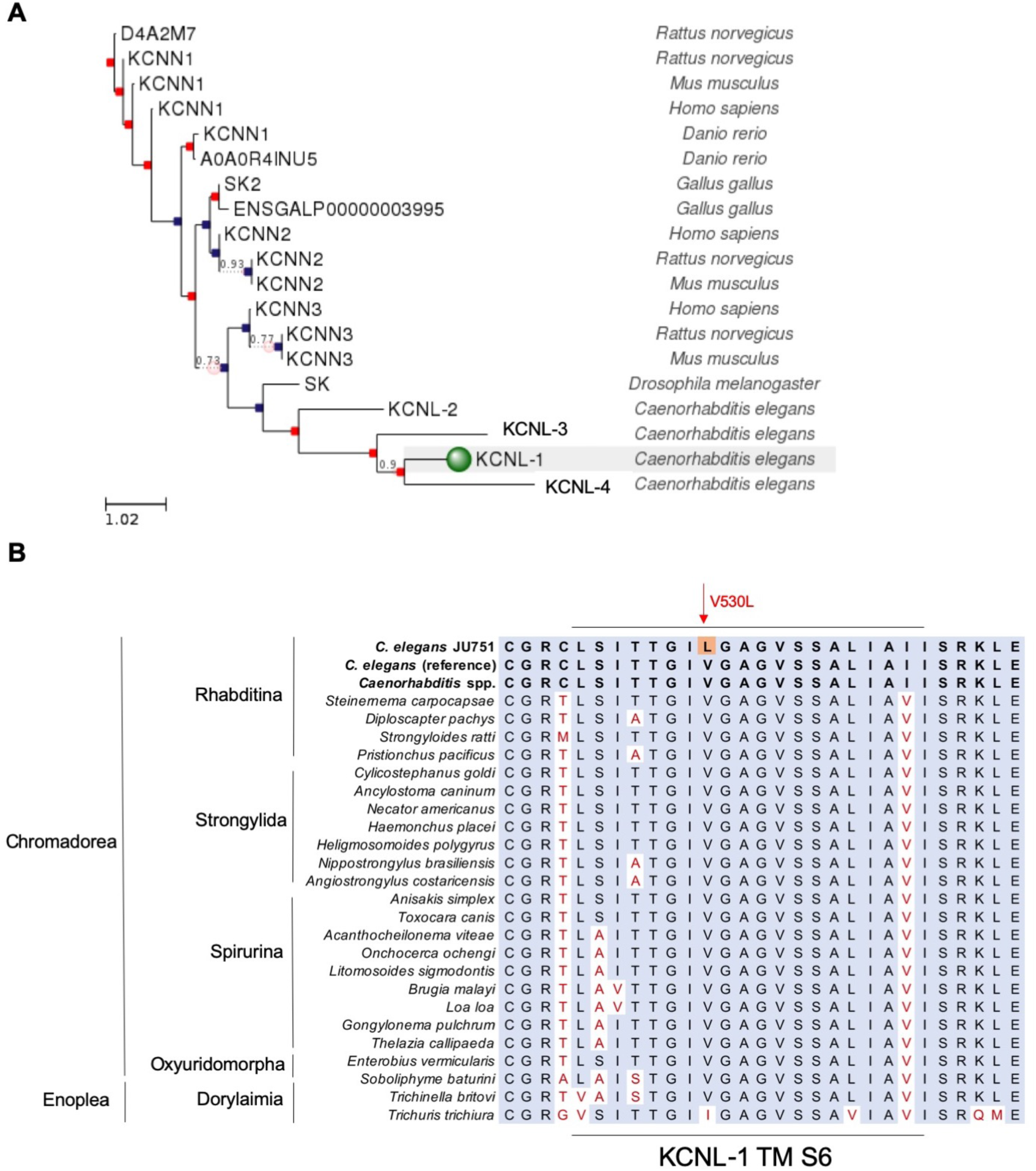
Phylogenetic relationships of KCNL-1 and conservation of the TM S6 region across nematodes. (**A**) Orthologs and phylogenetic relationships of the KCNL-1 protein (www.ensembl.org/Multi/Tools/Blast?db=core). *kcnl-1* is one of four *C. elegans* paralogues encoding small conductance calcium-activated potassium channel genes: *kcnl-1, kcnl-2, kncl-3, and kcnl-4. C. elegans* KCNL-1 is orthologous to several human genes, such as KCNN2 (potassium calcium-activated channel subfamily N member 2); KCNN3 (potassium calcium-activated channel subfamily N member 3); and KCNN4 (potassium calcium-activated channel subfamily N member 4). (**B**) Strong conservation of the KCNL-1 TM S6 region across the genus *Caenorhabditis* and distant nematode taxa.

**Fig. S9.**
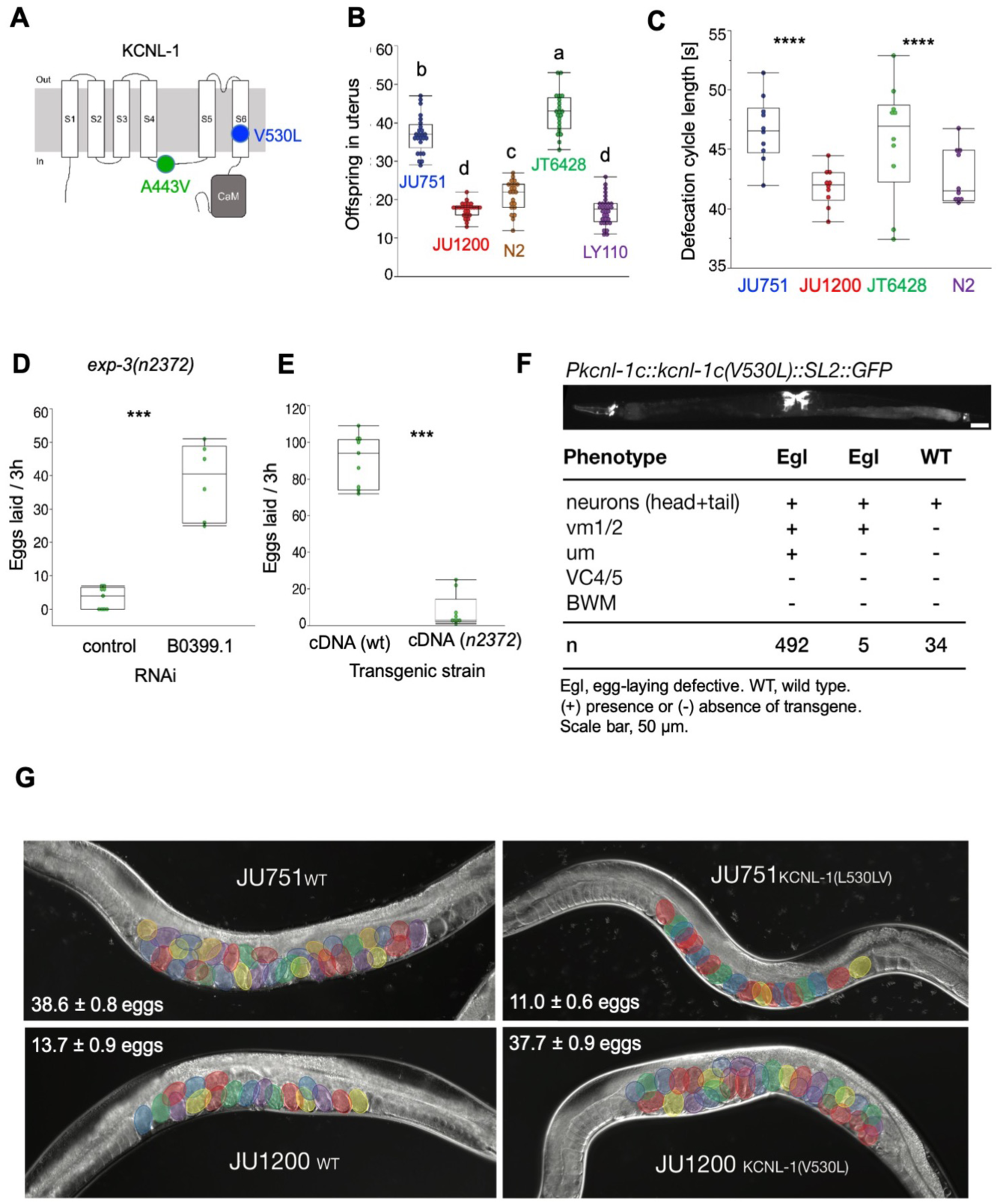
Characterization of *kcnl-1* mutations and *kcnl-1* expression. (**A**) Schematic view of the KCNL-1 ion channel subunit indicating amino acid changes in JU751 (V530L) and in the *exp-3(n2372)* mutant (A443V). V530L locates to the S6 transmembrane segment and A443V falls into the intracellular loop that connects the S4 and S5 transmembrane segment. (**B**) Offspring number *in utero* (embryos and larvae) of adult hermaphrodites (L4+24h) in the gain-of-function mutant *exp-3(n2372)* (JT6428), the reduction-of-function mutant *B0399.1(nf110)* (LY110), compared to reference wild-type strain N2 and wild isolates JU751 and JU1200. N=21-32/strain. (ANOVA, Tukey’s HSD: Values with the same letter are not significantly different from each other (P <0.05). (**C**) Defecation cycle length (hermaphrodites at the L4+24h stage) is significantly longer in JU751 compared to JU1200 (Nested ANOVA *strain(individual)*, P<0.0001) and in *exp-3(n2372)* (JT6428) compared to wild-type N2 (Nested ANOVA *strain(individual)*, P<0.0001). Each data point shown reflects the mean of five cycle periods measured for one individual (N=10/strain). (**D**) Knockdown RNAi of the gene *kcnl-1* (clone *B0399.1*, ORFeome library) (Rual et al., 2004) in the mutant *exp-3(n2372)*. Young adults were fed either RNAi bacteria containing an empty pDEST vector (control) or the *B0399.1* clone. After three hours, adults were removed and offspring was counted three days later. The strong egg retention (Egl) phenotype was partially suppressed by *B0399.1*. N=6-9 replicates, 5 individuals per replicate. (Mann-Wilcoxon Test, \*\*\**P* < 0.001). (**E**) Overexpression using extrachromosomal arrays: wild-type cDNA and mutant cDNA *exp-3*(*n2372*) with the A443V mutation (isoform c) were expressed using the *Pkcnl-1c* promoter. Transgenic adults were allowed to lay eggs for three hours after which adults were removed and offspring was counted three days later. The A443V mutation caused strongly reduced egg-laying. N=9 replicates, 5 individuals per replicate (Wilcoxon Test, \*\*\**P* < 0.001). (**F**) Mosaic analysis: restricted expression of the KCNL-1 V530L mutant to vulval muscles by using a *kcnl-1* promoter fragment (*Pkcnl-1c::kcnl-1c(V530L)::SL2::GFP*), which drives expression in vulval muscles but not VC neurons. Animals expressing this transgene in vulval muscles were incapable of laying eggs (n=492+5), while mosaic animals lacking the transgene in vulval muscles could lay eggs normally (n=34). (**G**) DIC images of adult hermaphrodites (L4+48h) in JU751, JU1200 and corresponding KCNL-1 allelic replacement lines. Mean and standard error of egg retention are shown for each strain (dataset from Fig. 3E).

**Fig. S10.**
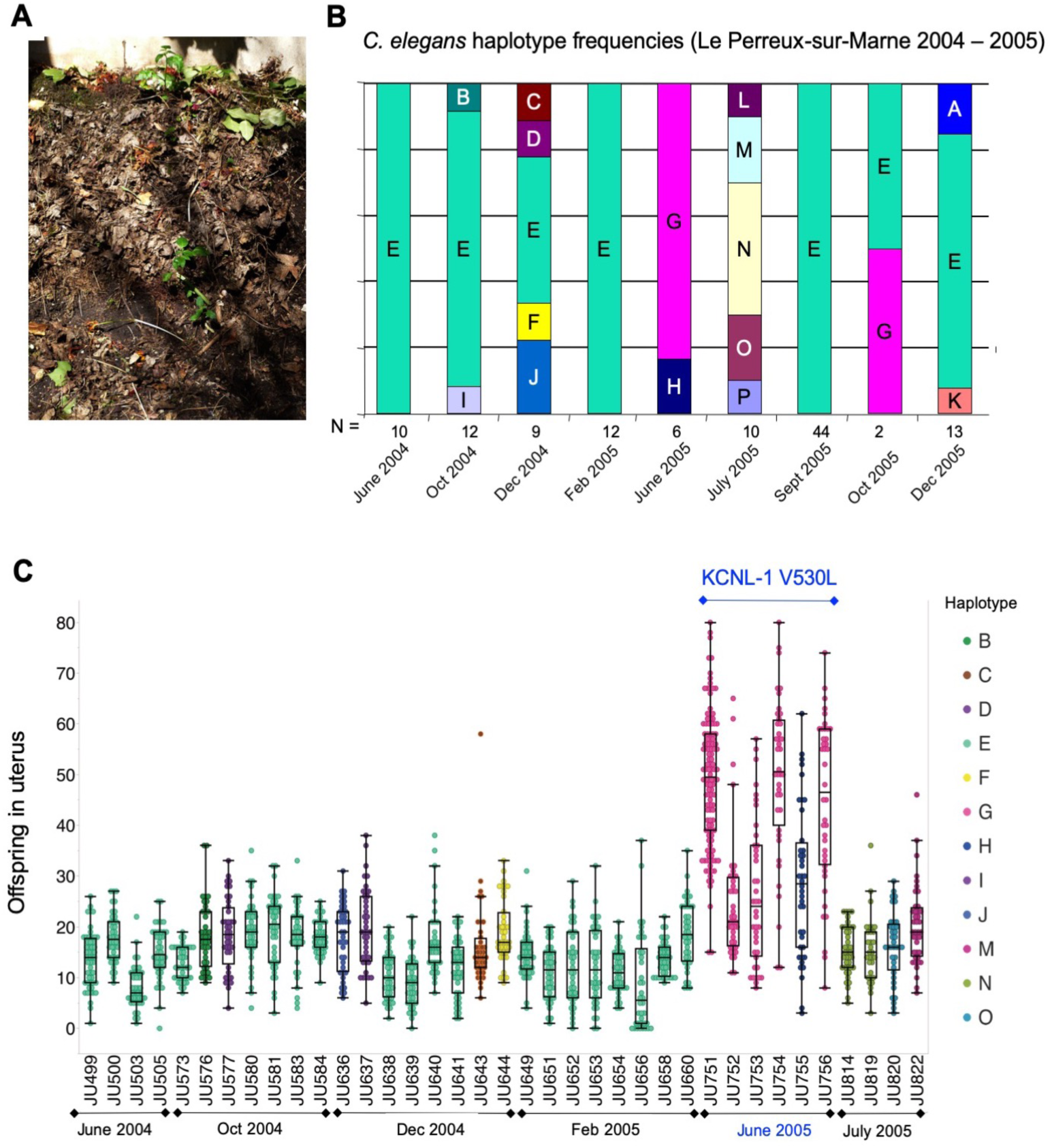
Natural origin of KCNL-1 V530L. (**A**) Photograph of the garden compost in Le Perreux-sur-Marne from which JU751 was isolated in 2005 (picture taken in 2018, courtesy of Jean-Antoine Lepesant). (**B**) Temporal dynamics of haplotype frequencies based on *C. elegans* sampling during 2004 and 2005 (Barrière and Félix, 2007). N indicates number of collected isolates. (**C**) Quantification of offspring *in utero* (embryos and larvae) of hermaphrodites (L4+48h) in select isolates (N=36) collected from the same compost heap in Le Perreux-sur-Marne throughout 2004 and 2005 N=31-120 individuals/per isolate. In addition to JU751, the five isolates (JU752, JU753, JU754, JU755, JU756) exhibited the KCNL-1 V530L polymorphism while none of the other isolates did (see also Data S1). These isolates, all collected on the same day (Barrière and Félix, 2007), showed significantly increased egg retention (and matricidal hatching). Note that isolates were established within a few hours after sample collection. All isolates collected between September and December 2005 were not cryopreserved and could thus not be phenotyped.

**Fig. S11.**
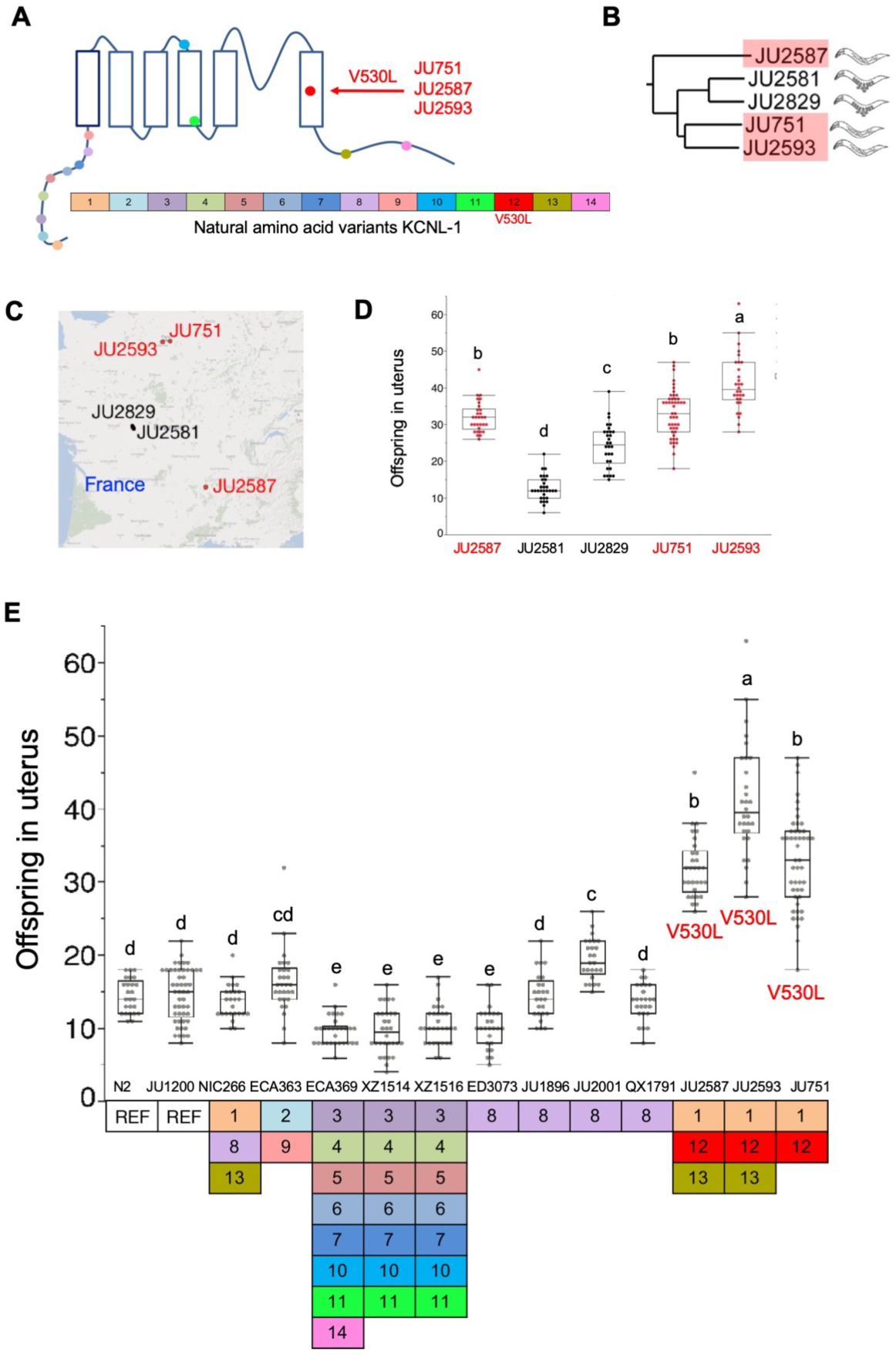
Survey of *C. elegans* natural variation in the KCNL-1. (**A**) Overview of amino acid variants in KCNL-1 present in one or more of 249 *C. elegans* wild isolates (Cook et al., 2017). 13 additional amino acid variants were detected in various other wild isolates at highly variable frequencies (see also Data S1). Approximate locations of the different variants in KCNL-1 are indicated in different colors. In addition to JU751, two other wild isolates (JU2587, JU2593) exhibit the KCNL-1 V530L variant. (**B**) The three isolates with KCNL-1 V530L are closely related based on whole-genome sequence similarity (Cook et al., 2017), indicative of a likely single origin of KCNL-1 V530L. Isolates JU2581 and JU2829 group with these strains but do not exhibit the KCNL-1 V530L variant. (**C**) Geographic distribution (limited to France) of the three isolates possessing the KCNL-1 V530L variant and the closely related strains JU2581 and JU2829. (**D**) Offspring number (embryos and larvae) *in utero* of closely related isolates, three of which showing the KCNL-1 V530L variant (red). N=30-47/strain. (ANOVA, Tukey’s HSD: Values with the same letter are not significantly different from each other (P <0.05). (**E**) Offspring number (embryos and larvae) *in utero* of wild isolates (hermaphrodites at L4+48h) carrying different KCNL-1 amino acid variants (indicated in colored boxes, REF: reference genotype), assessed on NGM plates in *ad libitum* food conditions. The three isolates with the KCNL-1 V530L variant (JU751, JU2587, JU2593) showed significantly increased egg retention and constitutive matricidal hatching. N=25-47/strain. (ANOVA, Tukey’s HSD: Values with the same letter are not significantly different from each other (P <0.05).

**Fig. S12.**
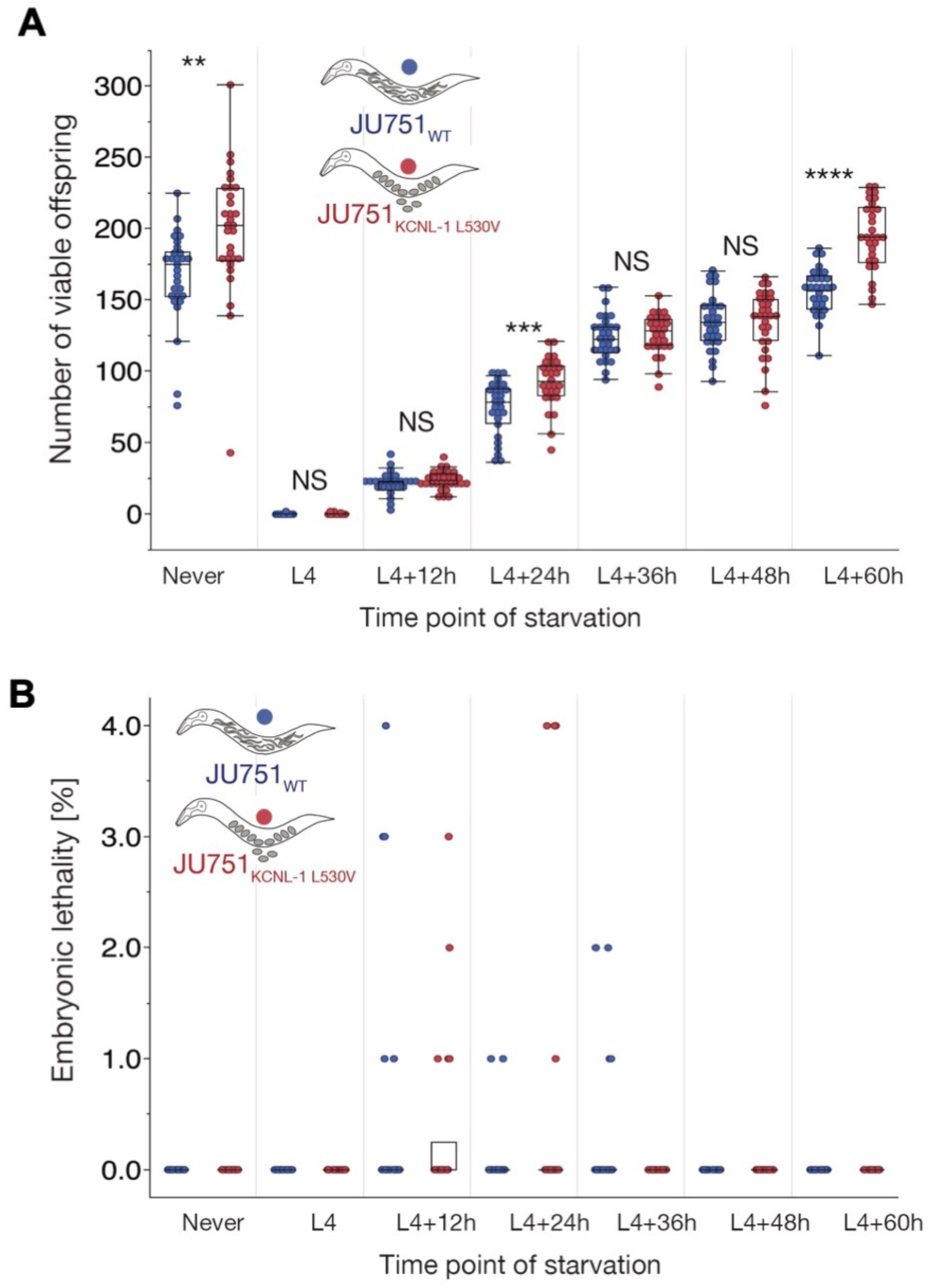
Fitness consequences KCNL-1 V530L in variable environments. (**A**) Lifetime production of viable offspring in selfing hermaphrodites in response to starvation encountered at varying maternal age: JU751_WT_ versus JU751 ARL_KCNL-1 L530V_. Age-synchronized populations were transferred from food (solid) to starvation (liquid) environment at different time points of development (N=27-30/strain/time point). Offspring number reflects combined viable larval offspring produced in food and starvation environments. (ANOVA separately performed for each time point: NS, not significant, \**P* < 0.05, \*\**P* < 0.01, \*\*\**P* < 0.001, \*\*\*\**P* < 0.0001). (**B**) Embryonic lethality in response to starvation encountered at varying maternal age: JU751_WT_ versus JU751 ARL_KCNL-1 L530V_ (from same experiment as in Fig. S12A, N=27-30/strain/time point). Embryonic lethality was very low for both strains across all time points (N=20 dead embryos in JU751_WT_, N=23 dead embryos in JU751 ARL_KCNL-1 L530V_). No embryonic lethality was observed in food conditions.

**Fig. S13.**
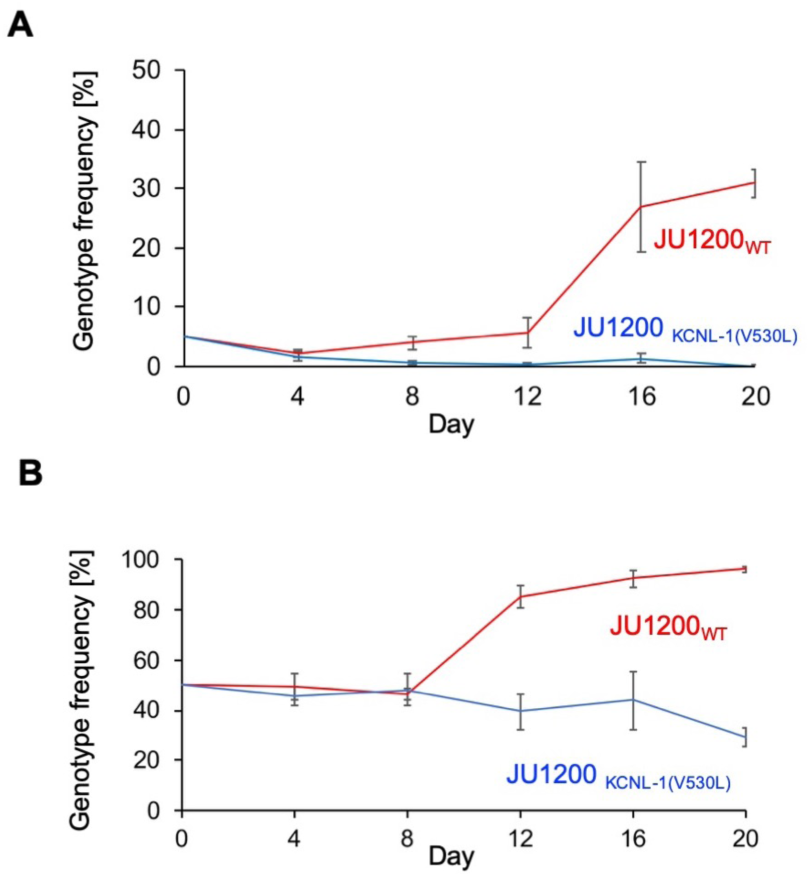
Competition and invasion experiments in constant food environments. (**A**) Competition of JU1200_WT_ and JU1200 ARL_KCNL-1 V530L_ against a GFP-tester strain (*myo-2::*gfp) with a starting frequency of 5% (invasion experiment). (**B**) Competition of JU1200_WT_ and JU1200 ARL_KCNL-1 V530L_ against a GFP-tester strain (*myo-2::*gfp) with a starting frequency of 50%.

### SUPPLEMENTARY TABLES

**Table S1.**
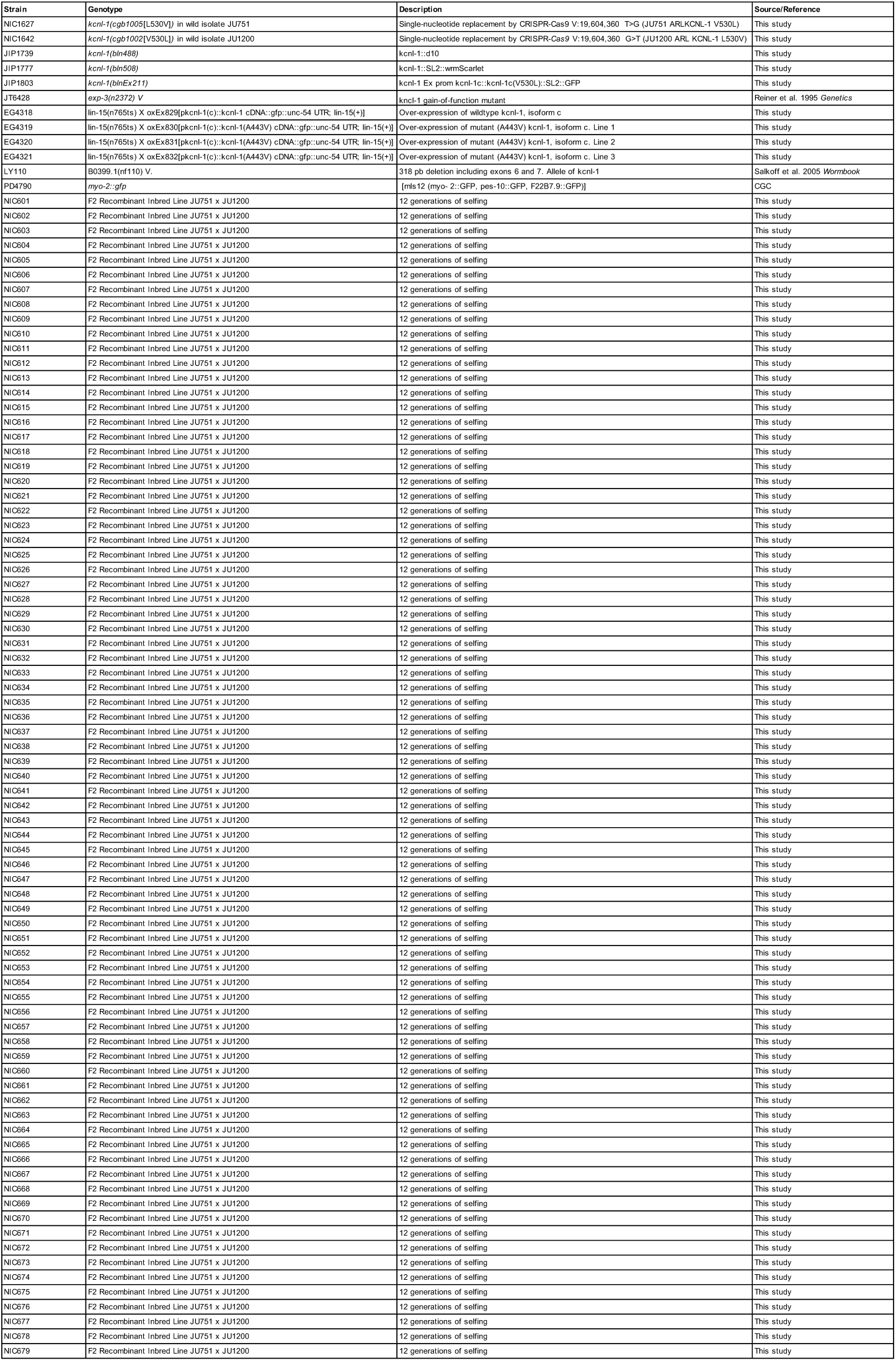
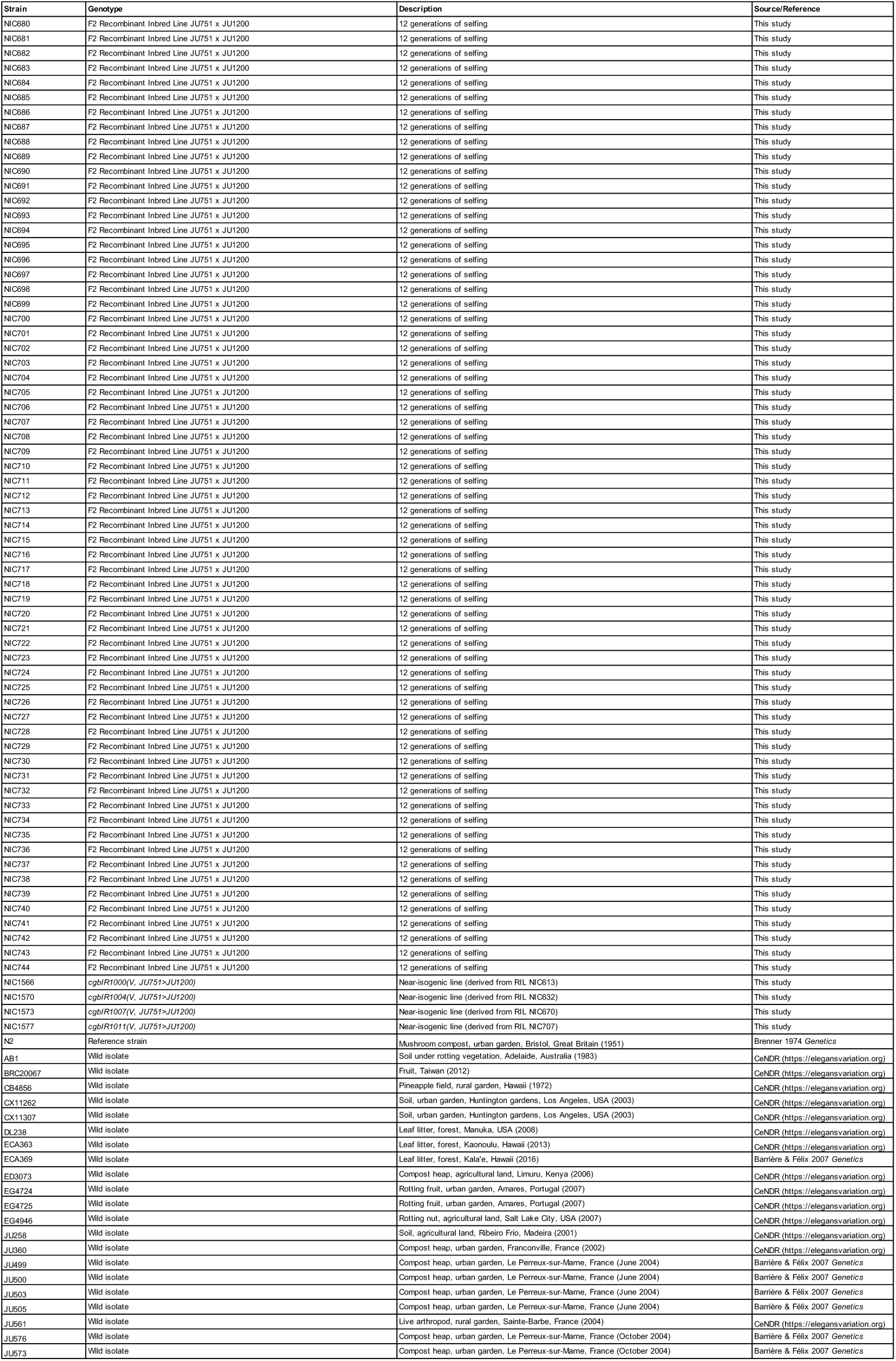
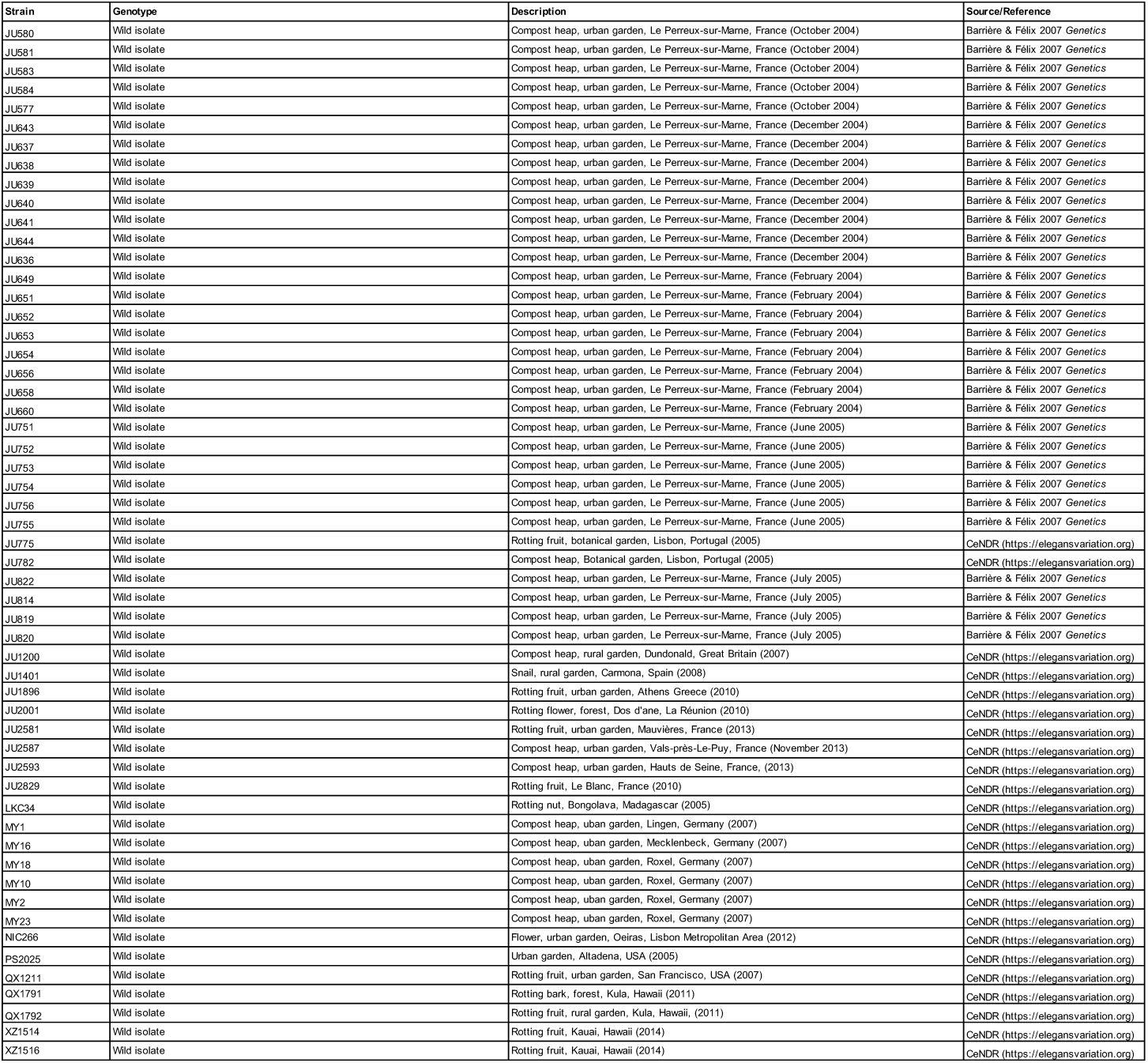
List of *C. elegans* strains used in this study.

**Table S2. (separate file)**

List of reagents used in this study.

**Data S1. (separate file)**

### Raw data in Excel format. Numbers below refer to different worksheets in Excel file

1. Natural variation in the plasticity of matricidal hatching across genetically divergent *C. elegans* wild isolates quantified in food (solid) and starvation (liquid) culture (Fig. 1A).
2. Temporal dynamics of matricidal hatching in JU751 versus JU1200: cumulative percentage of individuals containing one or more internally hatched larvae (Fig. 1C and S1C-E).
3. Egg retention and internal hatching in JU751 and JU1200 in *ad libitum* food conditions at 20°C (Fig. 1D-F and S1B).
4. Survival of JU751 and JU1200 hermaphrodites in *ad libitum* food conditions at 15°C and 20°C (Fig. 1G and S3A-B).
5. Effects of exogenous application of serotonin (5-HT) on egg-laying in JU751, JU1200 and the reference strain N2 (Fig. 1H)
6. Lifetime offspring production of selfing JU751 and JU1200 hermaphrodites (20°C) (Fig. S3C).
7. Sperm number of selfing JU751 and JU1200 hermaphrodites (20°C) (Fig. S3D).
8. Body size (perimeter) of JU751 and JU1200 hermaphrodites at the mid-L4 stage (20°C) (Fig. S3E).
9. Embryo size of JU751 and JU1200 hermaphrodites (20°C) (Fig. S3F).
10. Quantification of egg-laying behavior in JU751 and JU1200 (Fig. S4).
11. Genotypes of the 144 constructed JU751 x JU1200 F2 Recombinant Inbred Lines (RILs) (Fig. S5 and S6).
12. Raw phenotyping data (matricide) of JU751 x JU1200 F2 Recombinant Inbred Lines (RILs) (Fig. 2B and S5).
13. Simplified phenotyping data (matricide) of JU751 x JU1200 F2 Recombinant Inbred Lines (RILs) used for QTL mapping. (Fig. 2C and S5).
14. Phenotyping data (matricide) for near-isogenic lines (Fig. S7A).
15. Genotype data for near-isogenic lines (including data for relevant RILs and parental isolates) (Fig. S7B).
16. Offspring number *in utero* (embryos and larvae) of adult hermaphrodites (L4+24h) in the gain-of-function mutant *exp-3(n2372)* (JT6428), the reduction-of-function mutant *B0399.1(nf110)* (LY110), compared to reference wild-type strain N2 and wild isolates JU751 and JU1200 (Fig. S9B).
17. Defecation cycle length (hermaphrodites at the mid-L4+24h stage) of JU751 versus JU1200 and JT6428 versus N2 (Fig. S9C).
18. Knockdown RNAi of the gene *kcnl-1* (clone *B0399.1*, ORFeome library) in the mutant *exp-3(n2372)* (Fig. S9D).
19. Overexpression using extrachromosomal arrays: wild-type cDNA and mutant cDNA *exp-3*(*n2372*) with the A443V mutation (isoform c) were expressed using the *Pkcnl-1c* promoter (Fig. S9E).
20. Offspring number in uterus (embryos and larvae) of adult hermaphrodites (L4+48h) in JU751_WT_, JU751 ARL_KCNL-1 L530V_ and JU1200_WT_, JU1200 ARL_KCNL-1 V530L._ N=30/strain (Fig. 3E).
21. Temporal dynamics of matricidal hatching in JU751_WT_, JU751 ARL_KCNL-1 L530V_ and JU1200_WT_, JU1200 ARL_KCNL-1 V530L_ across the first three days of adulthood (L4+24h, L4+48h, L4+72h) (Fig. 3F).
22. Offspring number in uterus (embryos and larvae) of adult hermaphrodites (L4+36h) in JU751_WT_, JU751 ARL_KCNL-1 L530V_ and JU1200_WT_, JU1200 ARL_KCNL-1 V530L_ in food (solid) versus starvation (liquid). N=27-30/strain (Fig. 3G).
23. Lifetime offspring production of selfing hermaphrodites in JU751_WT_, JU751 ARL_KCNL-_ 1 L530V and JU1200_WT_, JU1200 ARL_KCNL-1 V530L_. N=43-47/strain (Fig. 3H).
24. Lifespan of selfing hermaphrodites in JU751_WT_, JU751 ARL_KCNL-1 L530V_ and JU1200_WT_, JU1200 ARL_KCNL-1 V530L_. N=29-45/strain (Fig. 3I).
25. Quantification of offspring *in utero* (embryos and larvae) of hermaphrodites (mid-L4+48h) in isolates (N=36) collected from the same compost heap in Le Perreux-sur-Marne throughout 2004 and 2005 (Fig. S10C).
26. Amino acid variants in KCNL-1 present in one or more of 249 *C. elegans* wild isolates (Cook et al., 2017). In addition to KCNL-1 (V530L), 13 additional amino acid variants were detected in various other wild isolates at highly variable frequencies (Fig. S11A).
27. Quantification of offspring *in utero* (embryos and larvae) of hermaphrodites (mid-L4+48h) in closely related isolates JU751, JU2581, JU2829, JU2587, JU2593 (Fig. S11D).
28. Offspring number *in utero* of wild isolates (hermaphrodites at L4+48h) carrying different KCNL-1 amino acid variants, assessed on NGM plates in *ad libitum* food conditions. (Fig. S11E).
29. Lifetime production of viable offspring in selfing hermaphrodites in response to starvation encountered at varying maternal age: JU1200_WT_ versus JU1200 ARL_KCNL-1_ V530L (Fig. 4A).
30. Embryonic lethality in response to starvation encountered at varying maternal age: JU1200_WT_ versus JU1200 ARL_KCNL-1 V530L_ (Fig. 4B).
31. Lifetime production of viable offspring in selfing hermaphrodites in response to starvation encountered at varying maternal age: JU751_WT_ versus JU751 ARL_KCNL-1 L530V_ (Fig. S12A).
32. Embryonic lethality in response to starvation encountered at varying maternal age: JU751_WT_ versus JU751 ARL_KCNL-1 L530V_ (Fig. S12B).
33. Competition of JU1200_WT_ and JU1200 ARL_KCNL-1 V530L_ against a GFP-tester strain (*myo-2::*gfp) with a starting frequency of 5% (invasion experiment) (Fig. S13A).
34. Competition of JU1200_WT_ and JU1200 ARL_KCNL-1 V530L_ against a GFP-tester strain (*myo-2::*gfp) with a starting frequency of 50% (Fig. S13B).
35. Invasive capacity of JU1200 ARL_KCNL-1(V530L)_ into a JU1200_WT_ population at a starting frequency of 5% across ∼15 generations (Fig. 4C).
36. Direct competition of JU1200_WT_ versus JU1200 ARL_KCNL-1 V530L_ at an initial 1:1 ratio across ∼15 generations (Fig. 4D).

